# Accurate sequence-dependent coarse-grained model for conformational and elastic properties of double-stranded DNA

**DOI:** 10.1101/2021.12.02.470889

**Authors:** Salvatore Assenza, Rubén Pérez

## Abstract

We introduce MADna, a sequence-dependent coarse-grained model of double-stranded DNA (dsDNA), where each nucleotide is described by three beads localized at the sugar and base moieties, and at the phosphate group. The sequence dependence is included by considering a step-dependent parameterization of the bonded interactions, which are tuned in order to reproduce the values of key observables obtained from exhaustive atomistic simulations from literature. The predictions of the model are benchmarked against an independent set of all-atom simulations, showing that it captures with high fidelity the sequence dependence of conformational and elastic features beyond the single step considered in its formulation. A remarkably good agreement with experiments is found for both sequence-averaged and sequence-dependent conformational and elastic features, including the stretching and torsion moduli, the twist-stretch and twist-bend couplings, the persistence length and the helical pitch. Overall, for the inspected quantities, the model has a precision comparable to atomistic simulations, hence providing a reliable coarse-grained description for the rationalization of singlemolecule experiments and the study of cellular processes involving dsDNA. Owing to the simplicity of its formulation, MADna can be straightforwardly included in common simulation engines.

## Introduction

Sequence-dependent conformational and elastic properties of DNA are of the utmost importance for its regulation in vivo, as they directly affect DNA-protein interactions.^1^ The detailed shape of a DNA fragment and its deformability are indeed key determinants of protein recognition and binding.^1,2^ For instance, the unique conformational properties of A-tracts are known to affect nucleosomal organization,^3^ as well as DNA replication and recombination. ^4^ Analogously, TATA-box elements present a strong deviation from the canonical B-DNA conformation, which is exploited to enhance binding by the TATA-box binding protein. ^1^ Moreover, proteins continuously exert mechanical stress on bound DNA. For example, torsion regulates the activity of topoisomerases and polymerases,^5,6^ as well as playing a central role in chromatin remodelling.^7^ Also DNA stretching is relevant in vivo, e.g. in the action of recombinases^8^ or for site recognition in nucleosomes.^9^

Together with the practical implications of DNA elasticity in nanotechnological applications,^10,11^ this fundamental interest has prompted a conspicuous amount of experimental efforts devoted to the detailed characterization of DNA conformational ensemble and elasticity. For the study of conformers, classical crystallographic and NMR studies are nowadays being complemented by X-ray interferometry^12,13^ and by single-molecule imaging with Atomic Force Microscopy (AFM)^14,15^ or cryo-electron microscopy.^16^ Single-molecule techniques like AFM, optical tweezers and magnetic tweezers are employed to assess the elastic properties of long DNA molecules, providing stiffness values for various mechanical perturbation modes such as stretching,^17–21^ twisting,^20,22–27^ bending^17–19,21,28–30^ and the coupling between them.^20,31–35^ At short length scales, interpretation of experiments becomes cumbersome due to both theoretical and experimental challenges. Indeed, direct AFM imaging has provided conflicting results on the bendability of short DNA fragments.^36,37^ For other experimental approaches such as cyclization assays,^38–41^ the elastic parameters can be obtained only by interpreting the data within specific theoretical frameworks, and conflicting conclusions have been reported based on different physical assumptions.^12,42–44^

Molecular simulations provide a valuable mean to complement experimental studies, particularly in view of the discrepancies commented above. All-atom molecular dynamics has been successfully employed to capture DNA elastic and conformational features^45–49^ as well as their dependence on sequence. ^35,50–53^

An evident limit of atomistic simulations originates from the associated high computational cost, which puts severe boundaries on the length and time scales that can be simulated. The need to overcome this barrier has fostered the development of coarse-grained approaches, where one selects only the degrees of freedom relevant to the problem at hand. An elegant solution is implemented in the software packages cgDNA^54^ and MC-eNN,^55^ where the nucleotides are represented as rigid frames and the energy of the system is described by means of a stiffness matrix. This approach is extremely efficient from a computational perspective and enables assessing the conformational ensemble of long DNA chains. ^56^ However, an important drawback lies in the absence of an explicit description of the backbone and the related difficulty in interfacing this representation with external perturbations present in most systems of interest, e.g. mechanical stress, confinement or a binding protein. A good compromise between computational efficiency and modelling flexibility is achieved by considering coarse-grained models in which effective particles represent the various moieties along the DNA molecule. ^57–65^ These models have been employed to describe a wide palette of systems involving DNA, including e.g. DNA origami,^66^ nucleosomes^67^ and cyclization 44 assays. ^44^

Despite this success, the coarse-grained models of the latter kind present in the literature do not satisfactorily capture all the main elastic features of double-stranded DNA at once (see e.g.^62,68^). In this work, we fill this gap by introducing the Mechanically-Accurate DNA (MADna) model, a sequence-dependent coarse-grained model whose parameters are tuned to reproduce local conformational features of double-stranded DNA and the stiffness of the various mechanical perturbation modes obtained via all-atom molecular dynamics. ^52^ We show that the model satisfactorily reproduces the atomistic values also for DNA fragments different than the ones employed for fitting, as well as experimental data from literature. Particularly, and at variance with existing models, our coarse-grained description captures the negative twist-stretch coupling recently unveiled by single-molecule force spectroscopy^31,32^ as well as the experimentally-determined sequence dependence of dsDNA helical pitch and persistence length. ^28^

The full details of MADna and the definitions of the various physical and geometrical quantities are provided in Materials and Methods. Nevertheless, they are shortly introduced also in Results and Discussion, making this section somewhat self-standing. In this way, the main findings of this work are presented succintly, leaving the interested reader to look up the technical details within the Material and Methods section.

## Methods

### Coarse-grained model

MADna describes a molecule of double-stranded DNA (dsDNA) by considering three effective particles per nucleotide, located in the geometric centres of the phosphate group, the sugar and the base, as proposed in the past in the 3SPN^57^ and TIS ^65^ models. In Fig.1, two cartoons are reported to compare the atomistic and coarse-grained description of a representative sequence. Further details on the coarse-graining procedure are reported in section S1 in the Supplementary Information. In spite of the similar level of coarse graining, MADna diverges from the other models in terms of the choice of bonded interactions both in the way they are built and in their parameterization. Particularly, our choice of bonded interactions enables capturing the structural heterogeneity predicted by atomistic simulations, as will be showed below when we comment on the results in Fig.2 and Fig.4i. On the other hand, we chose to fix the double-stranded structure by means of bonded interactions, which simplifies the implementation of the model (for instance MADna does not need modulation factors to account for anisotropic interactions) but makes it less general than the other ones as it cannot account for DNA thermodynamics. We will add the possibility of strand separation in future versions of MADna. As for the parameterization, MADna extracts information from a large dataset provided by atomistic simulations, while both TIS and 3SPN are parameterized by means of experiments. Experimental results are of course the ultimate source of information, but we decided to tune the parameters of MADna from atomistic simulations based on both the far richer amount of available information and their demonstrated accuracy in capturing local conformational and elastic features of dsDNA. As we show in the Results section, this choice is rewarded with a very good performance of MADna in capturing many distinct experimental observations on structure and mechanical response of dsDNA.

**Figure 1:**
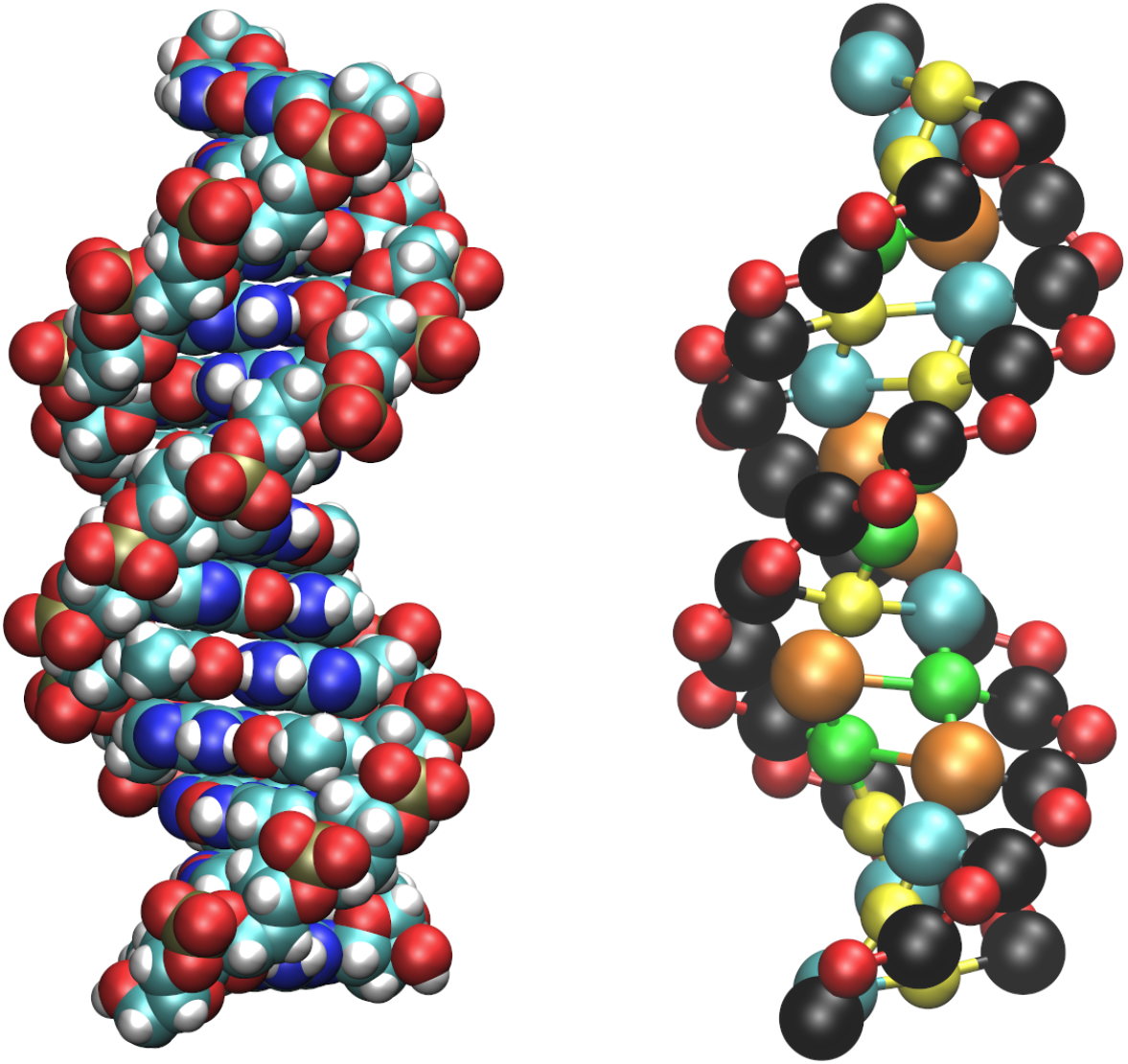
Atomistic (left) and coarse-grained (right) description for a representative dsDNA molecule with leading sequence 5’-CGCTACTTCGAGG-3’ in the B-DNA form as obtained by employing the NAB software.^69^ In the coarse-grained cartoon, the color code is the following: sugar ↔ black; phosphate group ↔ red; adenine ↔ green; cytosine ↔ cyan; guanine ↔ yellow; thymine ↔ orange. The size of each bead is proportional to the WCA radius of the corresponding moiety.

**Figure 2:**
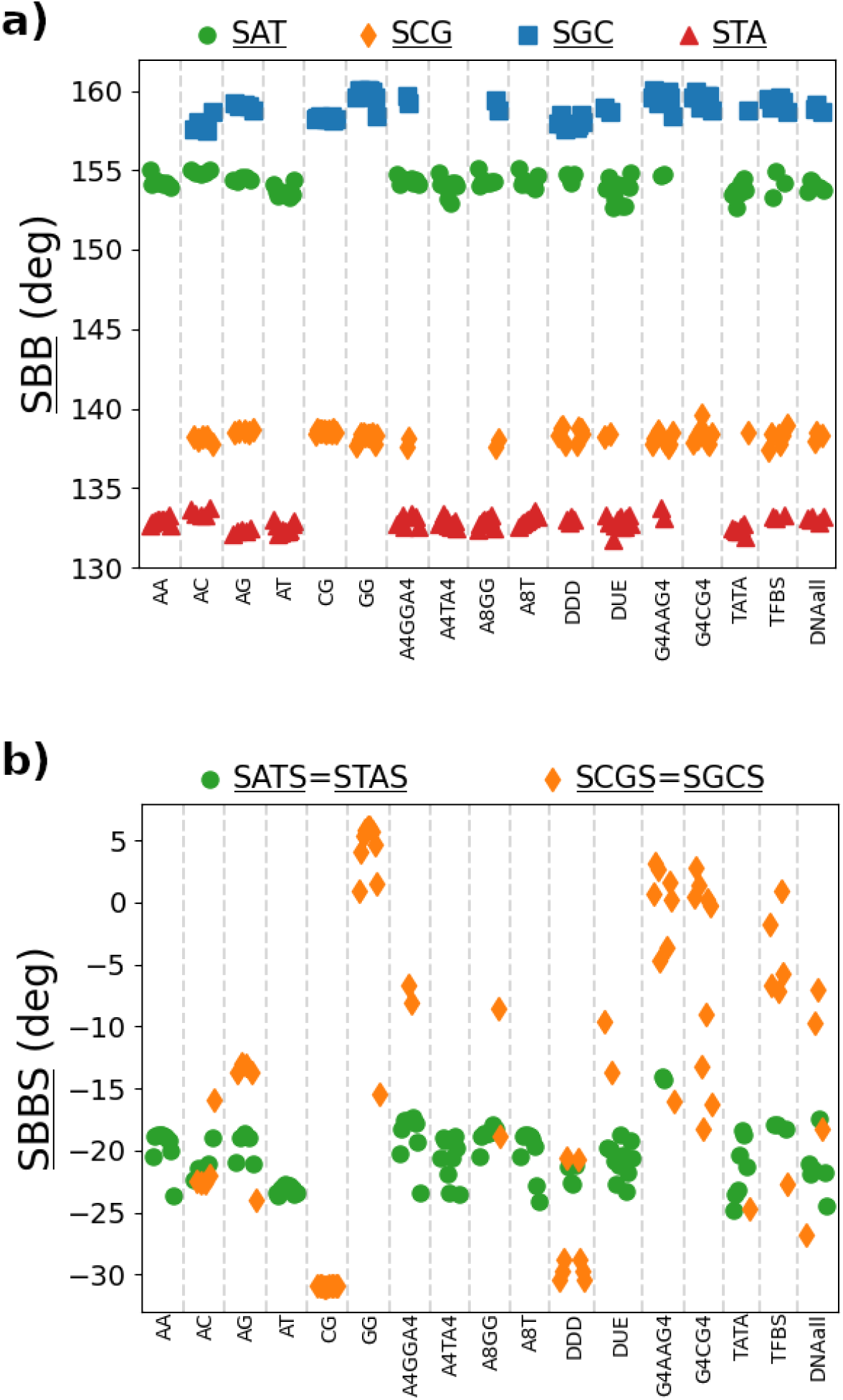
Average values obtained by coarse-graining the atomistic simulations for the angles <u>SBB</u> (a) and the dihedrals <u>SBBS</u> (b). Vertical lines separate the distinct sequences, which are listed in Section S3.1 in the Supplementary Information. In the legends, the various labels correspond to the particular bases involved in the local conformation under consideration. For instance, <u>SAT</u> considers a <u>SBB</u> angle in which an adenine is bound to the sugar. Note also that the dihedrals are symmetric with respect to the inversion of the involved bases, as it just corresponds to changing the arbitrary reference strand.

#### Bonded interactions

The double-stranded topology is fixed once and for all, and is maintained by introducing several two-, three- and four-body bonded interactions (from here on, we will refer to them as bonds, angles and dihedrals, respectively), which connect beads within the same strand as well as providing interstrand links.

Before going into the details of the bonded interactions considered, we first proceed to clarify the nomenclature used in this section. In order to label the bonded interactions, we indicate the sugar, phosphate and base beads as S, P and B, respectively. In this way, a bond between e.g. a sugar and a base is indicated as SB. Whenever the bonded interaction runs along the 5’-3’ direction of a strand, we add a corresponding tag at the two ends of the label. For instance, a bond between a sugar and a phosphate in the 5’-3’ direction is indicated as 5’-SP-3’. Analogously, we denote as 5’-SPSB-3’ a dihedral involving a sugar, a phosphate, the following sugar and the base attached to it (ordered according to the 5’-3’ direction of the strand). In order to keep in place the double-stranded structure, some bonded interactions involve both strands. Such bonded interactions are marked by underlining their label. For instance, the bonds accounting for Watson-Crick base pairing are indicated as BB-WC (in this specific case we also added “WC” in order to clearly distinguish these bonds from 5’-BB-3’, which account for stacking interactions). As another example, we denote as 5’-PSBB-3’ a dihedral involving in the 5’-3’ direction a phosphate, a sugar, the attached base and the base paired to it. We further note that, due to the asymmetry of the 5’-3’ direction, the order in which the same beads are considered is important. For instance, the bonds 5’-SP-3’ and 5’-PS-3’ are different from each other. This can be clearly seen by comparing the corresponding entries in Table S2 in the Supplementary Information, where we show the equilibrium values computed by coarse-graining the atomistic simulations used to parameterize MADna (see below). Analogously, the dihedrals 5’-SPSP-3’ and 5’-PSPS-3’ are different from each other, as can be seen from Table S4 in the Supplementary Information.

For the bonds potential *U*_bond_, a harmonic function was considered:

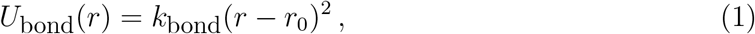

where *r* is the distance separating the two beads connected by the bond, *k*_bond_ is the elastic constant specific to the type of bond considered and *r*_0_ is the corresponding equilibrium distance. An analogous formula was used for the angle potential *U*_angle_:

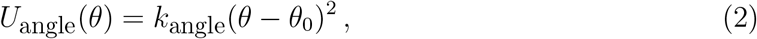

where *θ* is the angle characterizing the three-body bonded interaction, *k*_angle_ is the bending constant relative to the type of angle considered and *θ*_0_ is the equilibrium value of *θ*. Finally, the following formula was chosen for the dihedral potential *U*_dihedral_:

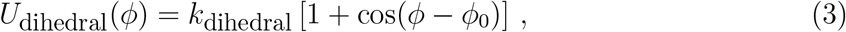

where *ϕ* is the dihedral angle, *k*_dihedral_ the elastic constant of the particular type of dihedral considered, and *ϕ*_0_ = *ϕ*_min_ – 180°, with *ϕ*_min_ being the equilibrium dihedral angle.

The general guideline in choosing how to build the topology was to obtain a minimal set of bonded interactions which reproduces the structural features obtained in all-atom simulations. With this spirit, we considered only bonded interactions involving up to a single step, and implemented the sequence dependence by setting different values of the parameters according to the particular step under consideration. For instance, if along the sequence we have a CT step, the bond 5’-BB-3’ accounting for the stacking interaction is implemented by means of Eq. (1), where *k*_bond_ and *r*_0_ are equal to the values assigned to a CT step. Analogously, the dihedral 5’-<u>PSBB</u>-3’ is implemented by means of Eq. (3), where kdih_e_d_ra_l and ϕ_0_ are set to the values corresponding to CT. The same is repeated for all the step-dependent bonded interactions considered, so that 16 different values are considered for the corresponding parameters depending on the involved step. Some local features are symmetric with respect to the 5’-3’ direction. The corresponding bonded interactions were then implemented as being dependent only on the corresponding base pair. For instance, for the bond SB we considered 4 possibilities (A,C,G,T), while for the bond <u>BB-WC</u> there were just two choices (AT or CG). Yet, this assumes that these local features do not depend on the neighboring steps, which is not always the case. As an example, we report in Fig.2 the averages obtained by coarse-graining the atomistic simulations for the angles <u>SBB</u> (Fig.2a) and the dihedrals <u>SBBS</u> (Fig.2b). The angles <u>SBB</u> are obtained by considering a sugar, the base attached to it and the corresponding Watson-Crick partner. For instance, we indicate as <u>SAT</u> the angle formed by a sugar, the adenine attached to it and the thymine paired with the adenine. As for the dihedrals <u>SBBS</u>, we similarly consider a Watson-Crick base pair and the corresponding sugars. In Fig.2, the points are color-coded according to a classification based on a single base pair. Hence, four possible choices are present for <u>SBB</u>, depending on the base being attached to the sugar (A,C,G,T), while only two choices are available for <u>SBBS</u>, corresponding to the Watson-Crick base pairs AT and CG. As shown in Fig.2a, the data for <u>SBB</u> nicely cluster around their average values, indicating that this angle is in practice independent of the rest of the sequence of the hosting DNA molecule and that it can thus be considered for bonded interactions depending only on the base pair. In contrast, the dihedrals <u>SBBS</u> show a wide variability, particularly in the case of CG base pairs (Fig.2b). This demonstrates that these dihedrals depend on their neighborhood, and are thus not apt for implementing bonded interactions depending only on the base pair. As we show in the Results, the heterogeneity of the SBBS dihedrals is quantitatively captured by MADna as an emergent property resulting from the interplay between the other bonded interactions. We also note that this is a distinguishing feature of MADna when compared to the TIS^65^ and 3SPN2^70^ models, where instead the base-pair dependent <u>SBBS</u> interactions are introduced as part of the model, thus preventing the possibility of capturing the heterogeneity suggested by the atomistic simulations.

The choice of the bonded interactions was obtained in practice by trial and error: starting from a far too simplistic description, where the beads were barely kept together by a minimal set of bonds, we incrementally added bonded interactions. At each iteration, we run simulations with the coarse-grained model and computed the averages obtained for local conformational features that were not yet directly implemented. We then compared this averages with the results obtained by coarse-graining the atomistic trajectories. If the comparison was not good, we then proceeded to add other bonded interactions. The iterative procedure stopped when we reached a satisfactory comparison. We also considered an opposite approach, where we started by considering far too many bonded interactions and deleted some of them until further eliminations resulted in disrupting the structure. The process has some arbitrarity in the properties being monitored, the order in which we add bonded interaction, etc. Yet, the high fidelity of the coarse-grained model in reproducing the atomistic values (as shown in the Results) validates the final choice of bonded parameters that we considered. The final set of employed bonds, angles and dihedrals is reported in Fig.3, together with some representative examples based on the dsDNA molecule sketched in Fig.1. Some of these bonded interactions are directly related to physically-meaningful features, such as hydrogen bonds between Watson-Crick pairs (labeled as <u>BB-WC</u> in Fig.3) and stacking interactions (5’-BB-3’ in Fig.3). Summing up, there are in total 202 distinct bonded interactions in the model. Further details are reported in Section S2 in the Supplementary Information.

**Figure 3:**
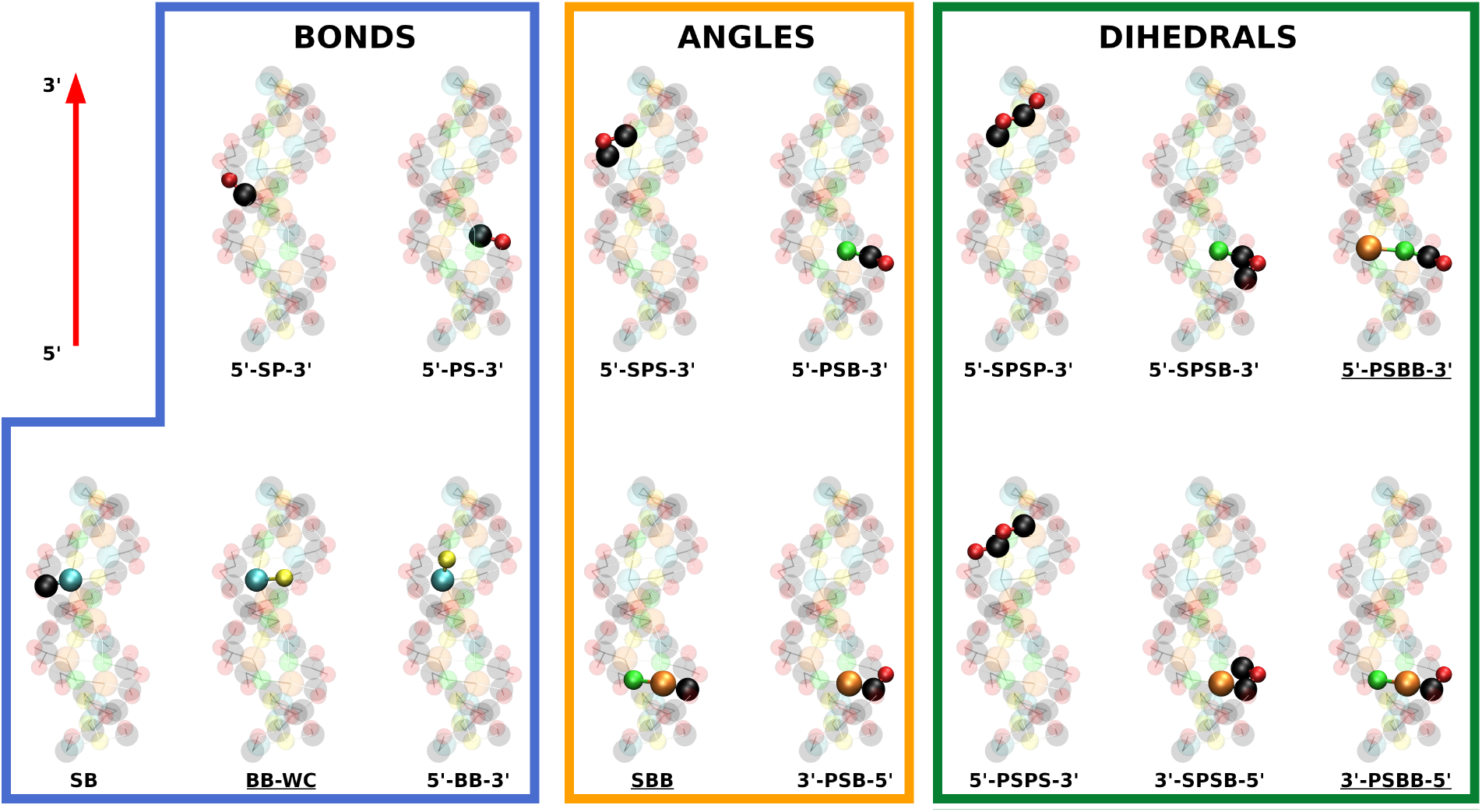
List of the various bonded interactions considered in the model, together with representative examples based on the same molecule as in Fig.1. Step-dependent bonded interactions are indicated by the presence of the tags 5’ and 3’ in their label. For interstrand interactions, the corresponding label is underlined. The letters present in the labels indicate sugars (S), phosphate groups (P) or generic bases (B). For clarity, all the selected examples involve beads belonging to the same strand, whose 5’-3’ direction is indicated by the arrow in the top-left panel.

Importantly, since all the bonded interactions involve at most two consecutive base pairs, any feature at scales larger than a single step is an emergent property originated from the propagation of the local interactions in combination with the electrostatic repulsion between phosphates (see below).

#### Excluded volume

Excluded-volume interactions were implemented by means of a Weeks-Chandler-Andersen (WCA) potential, i.e. by retaining the repulsive part of a Lennard-Jones interaction:

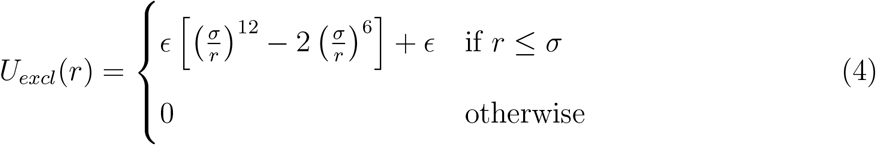

In the previous formula, *r* is the distance between the two particles, *ϵ* =1 Kcal/mol, while *σ* depends on the bead considered, as reported in Table S1 in the Supplementary Information.

#### Electrostatics

In order to account for the presence of charges along the backbone, a charge *q*·*e*_0_ was assigned to the beads corresponding to phosphate groups, with *e*_0_ = 1.6 · 10^-19^ C being the elementary charge. Although each phosphate carries a unit negative charge, in order to account for counterions condensation an effective reduced value *q* = −0.6 was considered.^65,70,71^ The salt-induced electrostatic screening was modeled via a Debye-Huckel interaction:^72^

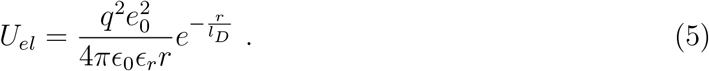

In the previous formula, *ϵ*_0_ = 8.859 · 10^−12^ F/m is the absolute permittivity; *ϵ_r_* = 78.3 is the relative permittivity of water; *l_D_* is the Debye length, defined as

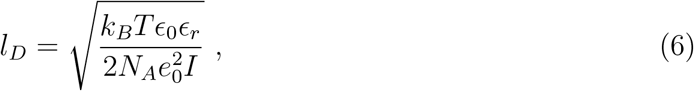

where *k_B_* = 1.38 · 10^-23^ J/K is Boltzmann’s constant; *T* is the temperature of the system in Kelvin; *N_A_* = 6.022 · 10^23^ mol^-1^ is Avogadro’s constant; *I* is the ionic strength of the solution in mM.

#### Determination of parameters

The parameters of the various sequence-dependent bonded interactions were tuned in order to reproduce the results of atomistic simulations from Ref.,^52^ which were performed in AMBER14^73^ with the parm99 force field^74^ including the bsc0 modifications.^75^ In those simulations, dsDNA molecules with different sequences were pulled under the action of forces ranging from 1 to 20 pN. Here, we used a subset of these simulations covering all possible base steps (we refer to this subset as “Learning sequences”, see Section S3.1 in the Supplementary Information) to determine the bonded parameters, while the rest of sequences were employed as test cases to benchmark the coarse grained model (“Testing sequences”). It has to be noted that, although parm99+bsc0 simulations reproduce well most structural features determined in experiments, ^76^ more precise modifications have been recently introduced, namely bsc1^77^ and OL15.^78^ Our reason to choose bsc0 was rooted in the convenience of having the atomistic data already available in our group,^46,52^ together with the reasonable precision of these modifications. ^76^ Future developments of MADna will include the reparameterization of the model via atomistic simulations based on bsc1 or OL15.

A first estimation of the parameters was obtained via Boltzmann Inversion of the all-atom simulations performed at 1 pN. In this regard, the atomistic trajectories were first coarsegrained according to the three-beads representation. Then, for each bond, angle and dihedral the ensemble averages were computed from the coarse-grained trajectories, which enabled fixing the values of *r*_0_, *θ*_0_ and *ϕ*_0_. Analogously, the elastic constants were determined in order to reproduce the size of fluctuations starting from Equations (1), (2) and (3). Further details on this procedure can be found in Section S2 in the Supplementary Information.

Coarse-grained simulations performed using the obtained force field showed that the set of parameters obtained by Boltzmann Inversion provides reasonable values for the elastic constants (see Fig. S1 in the Supplementary Information). Nevertheless, some of the parameters were further tuned in order to improve the quantitative agreement with the atomistic simulations, focusing on reproducing the elastic constants of the Learning Sequences (see Results and Discussion, and Section S2 in the Supplementary Information).

### Molecular Dynamics simulations

All the simulations were performed in LAMMPS (http://lammps.sandia.gov79). The temperature *T* = 300 K was maintained through a Langevin thermostat with damping constant *τ*_damp_ = 20 ps. The integration step was set to 20 fs.

#### Benchmark simulations

In order to benchmark the coarse-grained model, we performed pulling simulations of the Learning and Testing Sequences following the same protocol as in Refs. ^46,52^ A pulling force ***f*** was applied to the center of mass ***ξ***_2_ of the sugars belonging to the second base pair. Analogously, an opposite force –***f*** was applied to the center of mass ***ξ***_*l*_seq_–1_ of the second-to-last base pair (*l*_seq_ is the length of the sequence). The direction of the forces was taken along the line connecting ***ξ***_2_ and ***ξ***_*l*_seq_–1_. Simulations were performed for *f* =1, 5, 10, 15 and 20 pN. For each simulation, the system was initialized by considering the average structure created by the molecular builder (defined below) and the equations of motion were integrated for 20 ns. In order to ensure full equilibration, only the last 10 ns were considered for analysis. The convergence of a representative simulation is reported in Fig. S9 in the Supplementary Information. For each sequence and force, 100 independent simulations were performed. The ionic strength *I* was set at the same value as in the atomistic simulations from Refs. ^46,52^ and was computed as *I* = *N_ions_/N_A_V*, where *N*_ions_ is the number of counterions and *V* is the volume of the simulation box.

#### Persistence-length simulations

For the computation of the sequence-averaged persistence length, we considered 20 random sequences made of 100 base pairs. The sequences are listed in Section S3.2 in the Supplementary Information. For each realization, the molecule was initialized by considering the average structure obtained by the molecular builder (see below) and the simulation was run for 500 ns. In order to ensure full equilibration, the first 100 ns were discarded from the analysis. The convergence of a representative trajectory is reported in Fig. S10 in the Supplementary Information. For each sequence, 10 independent simulations were performed. The ionic strength was set at *I* = 150 mM.

In a second set of simulations, we studied the sequence dependence of persistence length and helical pitch, and compared our results with the experiments from Ref. ^28^ We considered 14 sequences of 100 base pairs obtained as the central fragments of the corresponding experimental ones.^28^ The sequences are listed in Section S3.2 in the Supplementary Information. The same protocol as for the sequence-averaged study was applied. The ionic strength was set at *I* = 1000 mM to ease the comparison with the predictions of CGDNA.^56^ The convergence of a representative trajectory is reported in Fig. S11 in the Supplementary Information. Simulations were performed with MADna and, for comparison, with oxDNA2^80^ (using the LAMMPS implementation^81^ and considering sequence-dependent stacking interactions) and the sequence-dependent model 3SPN2C. ^82^

#### Stretch-torsion simulations

In order to determine the elastic constants, a simulation setup was considered where a 40bp dsDNA molecule was subjected to the simultaneous action of a pulling force *f* and a torque *τ* directed along the z axis and with constant magnitude.

For each simulation, the molecule was initialized via the molecular builder and was aligned with the z axis. The two bottom base pairs and the corresponding sugars and phosphates were tethered to their initial position via harmonic constraints with constant 100 Kcal/mol Å^2^. A constant pulling force *f* oriented along the positive z direction was exerted on the center of the sugars belonging to the second base pair counting from the top. A harmonic constraint with constant 100 Kcal/mol Å^2^ was also applied to the x and y coordinates of the center to keep the molecule aligned with the z axis and avoid spurious effects originating from the overall orientational entropy.^83^ Finally, a constant torque *τ* was imposed on the two top base pairs and the corresponding sugars and phosphates.

Simulations were performed at *τ* = 0 for values of the force *f* = 2, 4, 6, 8, 10, 15, 20, 25, 30, 35, 40 pN, and at *f* = 2 pN for values of the torque *τ* = 0, 5, 10, 15, 20, 25, 30 pN·nm. To account for sequence-induced heterogeneity, five random sequences were generated, to which we refer as sequences ST1,…,ST5 (the sequences are listed in Section S3.3 in the Supplementary Information). Moreover, to check the dependence of the elastic constants on the length of the dsDNA molecule, we also performed simulations on shorter sequences containing 20 base pairs, to which we refer as sequences ST1-short,…,ST5-short (the sequences are listed in Section S3.3 in the Supplementary Information). For each combination of sequence, force and torque, 10 independent simulations of length 250 ns were run. The first 50 ns were not considered for analysis in order to let the system equilibrate. The convergence of a representative trajectory is reported in Fig. S12 in the Supplementary Information. Simulations were performed with MADna, oxDNA2 and 3SPN2C. In all cases, the ionic strength was set at 150 mM. The four base pairs at each end were discarded from the analysis.

Finally, in order to study the effect of sequence on the elasticity, we considered another set of simulations with phased A-tracts, which are listed in Section S3.3 in the Supplementary Information. The same protocol as for the rest of the simulations was applied.

#### Twist-bend coupling simulations

In order to determine the softer response to torque due to the coupling between twist and bending, we peformed simulations similar to the ones of the previous section. Here, 150bp-long sequences were put under the action of a pulling force directed along the z axis and acting on one end of the molecule. The other end was tethered with the same protocol as in the previous section. Note that the pulled end is here free to move in the x and y directions in order to let the molecule bend.

Simulations were performed for values of the force *f* in steps of 0.05 pN within the range 0.05 pN≤*f*≤ 0.50 pN, and in steps of 0.25 pN within the range 0.50 pN≤ *f* ≤ 2.50 pN. Three different sequences were considered, to which we refer as TB1,TB2,TB3 (the sequences are listed in Section S3.4 in the Supplementary Information). For each combination of sequence and force, three independent simulations of length 2000 ns were run. The first 1000 ns were not considered for analysis in order to let the system equilibrate. The convergence of a representative trajectory is reported in Fig. S13 in the Supplementary Information. Simulations were performed with MADna, 3SPN2C and oxDNA2.

### Definitions and protocols

#### Basic definitions

We will indicate as ***P***_*s,i*_, ***S***_*s,i*_ and ***B***_*s,i*_ the position vectors of the beads corresponding to the *i*-th phosphate group, sugar and base, respectively, and located on strand *s* = 1, 2. The index *i* is counted in the 5’3’ direction of the arbitrarily-chosen strand 1. Note that *i* = 1,…, *n* for sugar and base beads, while *i* = 2,…, *n* for the phosphate groups, where *n* is the total number of base pairs (compare Fig.1). Hence, there are in total *N* = 2[(*n*–1)+*n*+*n*] = 6*n*–2 beads, where the factor of 2 is included to account for the presence of the two strands. Handles were always discarded from the analysis, so the various indexes introduced here refer to the subfragment under consideration.

#### Crookedness

The crookedness *β* accounts for the deviation of a dsDNA molecule from a straight line^52^ (see Fig.S2 in the Supplementary Information). For short fragments (such as the Learning and Testing Sequences considered here), thermally-induced bending is negligible, hence in this case *β* effectively quantifies the spontaneous bending of the molecule. The definition of *β* is here modified with respect to its original formulation to adapt it to the coarse-grained context. The contour length *L* of the line connecting the base-pair centers along the molecule is 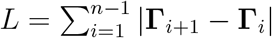, where **Γ**_*i*_ = (***B***_1,*i*_ + ***B***_2,*i*_)/2. The crookedness *β* is defined as

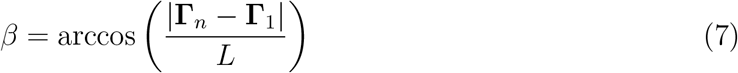

If the line connecting the centers is perfectly straight, then *β* = 0. The more curved the line, the larger the corresponding *β*. As a reference, we computed the crookedness for the conformations obtained via NAB^69^ for the Learning and Testing Sequences, obtaining an average value *β* ≃ 0.17.

#### Helical parameters

For a visual description of the definitions reported in the present section, see Fig.S2 in the Supplementary Information. Due to the limited information on local coordinates, the definition of the helical parameters has to be adapted to the coarse-grained framework. Particularly, the representation of the bases as single beads prevents defining the local reference frames, which are crucial for the standard definitions of helical parameters. ^84,85^ Taking advantage of the short length of the Learning and Testing Sequences, the helical axis 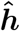 in this case was defined globally as the axis of the cylinder best fitting the position of phosphates in the double helix (see Section S4.1 in the Supplementary Information), which was also employed to estimate the diameter of DNA. In order to define the h-twist, we first introduce the sugar separation vector ***ζ***_*i*_ ≡ ***S***_2,*i*_ – ***S***_1,*i*_. The h-twist for the step *i,i* + 1 was defined by 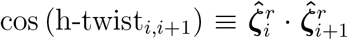, where 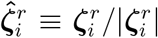 and 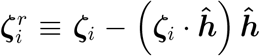 is the projection of ***ζ***_*i*_ onto the plane perpendicular to the helical axis. Following the usual convention, ^84^ the sign of the h-twist was determined according to the sign of the product 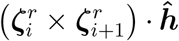. The employment of the positions of the sugars in the definition of the h-twist was motivated by the high correlation found with the standard definition in 3DNA^84^ (see Fig. S3 in the Supplementary Information). Finally, the h-rise for the step *i,i* + 1 was computed as 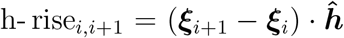, where ***ξ***_*i*_ ≡ (***S***_1,*i*_ + ***S***_2*i*_)/2 is the center of the sugars of base pair *i*.

#### Grooves

For a visual description of the definition of groove geometry, see Fig.S2 in the Supplementary Information. The quantification of the geometry of grooves was assessed in a similar way as in Curves+.^85^ The positions of the phosphate groups in each strand were interpolated via centripetal Catmull-Rom splines.^86^ Particularly, the helical fragment connecting ***P***_*s,i*_ and ***P***_*s,i*+1_ was interpolated by considering ***P***_*s,i*–1_, ***P***_*s,i*_, ***P***_*s,i*+1_, ***P***_*s,i*+2_ as control points. For each of the terminal phosphates, the missing external control point was built by extrapolation of the h-rise and h-twist of the last step. In order to determine the groove geometry corresponding to base pair *i,* we first considered the midpoint ***M***_1,*i*_ between ***P***_1,*i*_ and ***P***_1,*i*+1_ along the interpolated curve. Then, starting from the analogous midpoint ***M***_2,*i*_ on the second strand, we followed the corresponding interpolating curve in both directions while computing the distance from ***M***_1,*i*_, until a local minimum in the distance was first encountered in correspondence of two points denoted as ***F***_+,*i*_ and ***F***_−,*i*_. From the two identified minimal distances, we subtracted 0.58 nm to account for the van der Waals radius of the backbone^85^ and assigned the resulting values to the width of major and minor groove. In order to compute the depth of each groove, we considered the midpoints on the width vectors, i.e. 0.5(***M***_1*i*_ + ***F***_+,*i*_) and 0.5(***M***_1,*i*_ + ***F***_−,*i*_), and computed their distance from the base-pair center **Γ**_i_. Again, in order to account for the van der Waals radii, we estimated the final groove depths by subtracting 0.35 nm from the calculated distance. ^85^

### Determination of elastic constants

In order to interpret single-molecule experiments, dsDNA is often modelled as an elastic rod. When a pulling force *f* and a torque **τ** are exerted along the direction of the rod, the elastic energy *U* reads:^22,31^

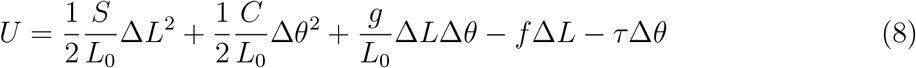

where *S* is the stretching modulus; *C* is the twist modulus; *L* is the extension; *θ* the twist in radians; Δ*L* ≡ *L* – *L*_0_ and Δ*θ* ≡ *θ* – *θ*_0_, with *L*_0_ and *θ*_0_ being the equilibrium extension and twist at zero force and torque. In the right-hand side of Equation (8), the first and second terms are stretching and twisting energy, respectively; the third term is the stretch-twist coupling energy; the fourth and fifth terms are the work performed by the external force and torque, respectively.

Simultaneous minimization with respect to extension and force (*∂U*/*∂*Δ*L* = 0 and *∂U*/*∂*Δ*θ* = 0) leads to the following equations for the elastic equilibrium:

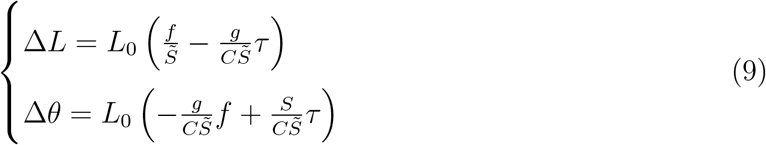

where 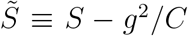 is the effective stretching modulus. From the previous formulas, the sign of *g* can be immediately ascertained by looking at the behavior of Δ*L* as a function of *τ* at constant force, with positive and negative correlation implying *g* < 0 and *g* > 0, respectively (an analogous response is obtained at constant *τ* for Δ*θ* as a function of *f*). For a quantitative assessment of the elastic constants, three equations are needed. Different strategies were pursued for the two simulation protocols considered in this work.

#### Benchmark simulations

In this case, the torsion angle is estimated as 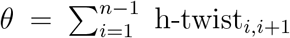. The extension is computed as *L* = |**Γ**_*n*_ – **Γ**_1_|. Moreover, no torque is applied, *τ* = 0. From Equation (9), one has Δ*L* = *A*_1_*f* and Δ*θ* = *A*_2_*f*, with 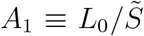 and 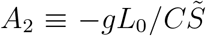. The third equation is provided by applying the equipartition theorem to Equation (8) and neglecting the twist-stretch coupling term. ^31^ This is not a problem for the present purposes, since it affects equally the analysis of both atomistic and coarse-grained results, thus not jeopardizing the quality of their comparison, although this detail should be kept in mind for the physical interpretation of the results. Application of the equipartition theorem gives on average Δ*θ*^2^ = *k_B_TL*_0_/*C*. Inverting these formulas, one finds

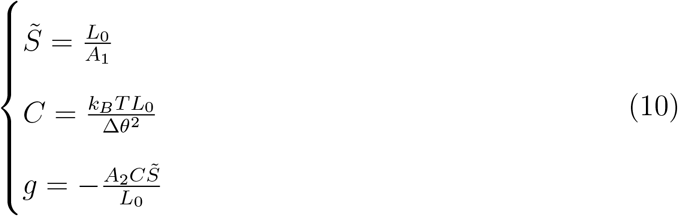

Fitting the linear dependence of *L* and *θ* on the pulling force *f*, we computed the constants *L*_0_, *θ*_0_, *A*_1_ and *A*_2_. Δ*θ*^2^ was instead calculated directly from the simulations performed at *f* = 1 pN. Plugging the values of these quantities into Equation (10), we finally obtained the values of 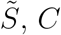 and *g* reported in Fig.5.

As for the crookedness constant *k_β_*, following Ref.^52^ we define it via the relation cos *β* = cos*β*_0_ (1 + *f*/*k_β_*), where *β*_0_ is the crookedness at zero force. *k_β_* is then suitably extracted from the slope of cos *β* versus *f*.

#### Stretch-torsion simulations

Taking advantage of the directionality imposed by the external force, the torsion angle in this case is computed as 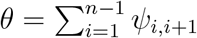, where the angle ψ is calculated as the h-twist but considering the rejection of ***ζ***_*i*_ from the direction of the force instead of the helical axis. In the same way as done above, for **τ** = 0 one has Δ*L* = *A*_1_*f*, with 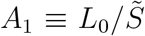. At variance with the Benchmark simulations, the presence of an imposed torque enables deriving all the equations without resorting to the analysis of the fluctuations. Indeed, for a constant pulling force *f* = *f*_0_, Δ*L* = constant + *A*_3_*τ* and Δ*θ* = constant + *A*_4_*τ*, with 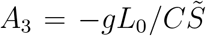 and 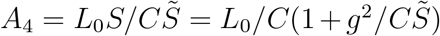. From the knowledge of the constants *A*_1_, *A*_3_, *A*_4_ one can obtain the elastic constants as

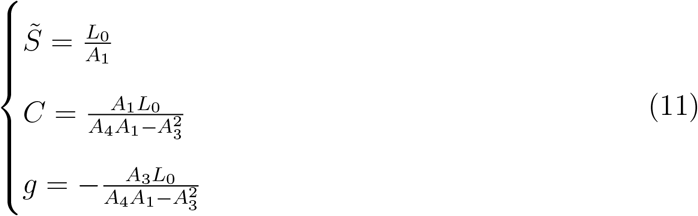

Fitting the linear dependence of Δ*L* on the pulling force *f* at *τ* = 0, we computed the constants *L*_0_ and *A*_1_. Analogously, fitting of Δ*L* and Δ*θ* versus *τ* at *f* = 2 pN enabled the computation of *A*_3_ and *A*_4_. Plugging the values of these quantities into Equation (11), we finally obtained the values of 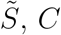 and *g* reported in Fig.6.

### Computation of persistence length

In order to compute the persistence length, for each base pair *i* we considered the geometrical center ***ξ***_*i*_ = (***S***_1,*i*_ + ***S***_2,*i*_)/2 of the sugar beads. By introducing the displacement vector ***R***_*ij*_ ≡ ***ξ*** – ***ξ***_*i*_, the *i*th tangent vector is defined as 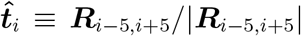. The contour length *l* separating the points of application of two consecutive tangent vectors is computed as the average modulus of the displacement vector. The overall correlation function *c_p_* is defined as

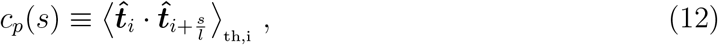

where *s* is the contour length, while 〈…〉_th,i_ denotes the average over both the conformational ensemble and the value chosen for *i*. Therefore, *c_p_* computes the decay of the orientation of the tangent vectors by taking thermal fluctuations into account. Analogously, the static correlation function *c_s_* is defined as

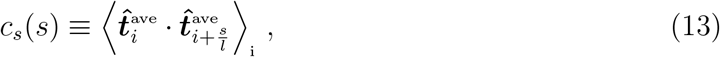

where 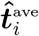 is the ith tangent vector computed on the average structure (obtained from the molecular builder) and 〈…〉_ti_ denotes the average over the value chosen for *i*. The static correlation function quantifies the decay of the orientation of tangent vectors along the average structure. Finally, following Ref.,^56^ the dynamic correlation function is defined as

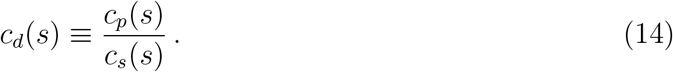

This is in fact equivalent to impose, for each sequence, that the harmonic relation 1/*l_p_* = 1/*l_s_* + 1/*l_d_* proposed by Trifonov and coworkers holds exactly.^56,87^

### Errors

Indeterminacies on the computed quantities were estimated according to the procedures below. Note that error bars are not present in the Figures whenever they are smaller than the size of symbols.

#### Observables

For all coarse grained simulations, several independent realizations were performed for each case, whose number depends on the particular set of simulations considered (see above). For each computed observable, the indeterminacy on the average was obtained as the standard error of the mean computed across the independent realizations. As for Δ*θ*^2^, the standard error of the variance was considered. In the case of the atomistic trajectories, the time series of each observable was decorrelated by performing a block analysis, from which the error was then estimated. ^88^

#### Elastic constants

In order to account for the errors on the elastic constants, the following procedure was employed. In all cases, the constants were obtained starting from a linear relation *y* = *ax* + *y*_0_, where *x* is either the force or the torque, while *y* is a certain observable (e.g. the extension). Let 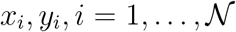 be the 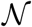 average values of the variables, and let *dy_i_* be the error associated to *y_i_* (*x_i_* does not have an associated error). Assuming the values of *y* to be independent and distributed normally, we iteratively considered sets of variables 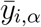 extracted from Gaussians with average *y_i_* and standard deviation *dy_i_*, where 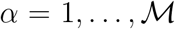 indicates the particular realization considered. For each such realization, we obtained the fitting parameters *a_α_* and *y*_0,*α*_ by linear regression. The mean slope was finally obtained as the average 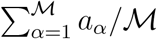, while the associated error was estimated as the corresponding standard deviation. We checked numerically that a satisfying convergence was obtained for 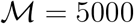.

### Molecular builder

The results obtained for the Learning Sequences were employed to devise a molecular builder, which provides sequence-dependent average structures to be used as initial configurations or for the computation of the static persistence length. For each step XY, with both X and Y chosen among the four bases A,C,G,T, we considered the simulations run for the Learning Sequence which contains it. For instance, the sequence PolyAA was considered for the steps AA and TT. The corresponding trajectories were aligned in order to minimize the root-mean-square deviation (rmsd) of the central step XY, computed according to the Kabsch algorithm.^89^ After this operation, the average coordinates of the ten beads belonging to the step (four sugars, four bases and two phosphates) were computed. These coordinates can be employed to build the average structure of a molecule, where the junctions between adjacent steps are aligned in order to minimize the rmsd of the overlapping base pair. As an example, let us consider the molecule with sequence 5’-ACT-3’. Two steps are present: AC and CT. The molecule is initiated by considering the average coordinates of the step AC. At this point, we note that the four beads (two sugars and two bases) belonging to the second base pair of the step AC are the same as the four beads belonging to the first base pair of the next step, CT. The average coordinates of the latter are then translated and rotated in order to minimize the rmsd between the two sets of coordinates for the overlapping beads. The final coordinates for the four overlapping beads are computed as the mean of the two aligned sets. This procedure can be iterated for molecules of arbitrary length. As a last step, the final structure is translated so that the origin corresponds to the first sugar bead and subsequently aligned to the z axis.

## Results and discussion

### Summary of coarse-grained model

In line with previous work, ^57,65^ the coarse grain considered in MADna describes each nucleotide by means of three beads, each centered on the sugar, phosphate group and base, respectively (Fig.1). The beads interact with each other via steric interactions and, in the case of phospate groups, electrostatic repulsion. The double-stranded molecular structure is maintained by introducing bonded interactions (Equations (1), (2), (3) and Fig.3), with parameters depending on the sequence up to the level of a single step. The parameters were determined by Boltzmann Inversion of atomistic trajectories from literature, ^52^ whose sequences encompass all the possible steps (hereby referred to as Learning Sequences, see section S3.1 in the Supplementary Information for details). Further tuning was performed in order to reproduce the elastic response to an external pulling force computed for the same set of sequences. Full details of the model and the coarse-graining procedure are found in the Methods and in Sections S1 and S2 in the Supplementary Information.

### MADna reproduces conformational features from atomistic simulations

In order to benchmark the optimized simulation setup, we proceeded with a systematic comparison between coarse-grained predictions and results from atomistic simulations. We performed coarse-grained simulations of dsDNA molecules under the action of a pulling force *f* ranging in the interval 1-20 pN, following the same protocol as in Refs.^46,52^ (see Methods for details on the implementation). Apart from the Learning Sequences, we considered a second, independent set of molecules (Testing Sequences), which has also been studied in Refs.^46,52^ The Testing Sequences are reported in Section S3.1 in the Supplementary Information and include biologically-relevant structures as well as synthetic A-tracts fragments. Both conformational and elastic properties were considered for the analysis.

The study of various conformational quantities is reported in Fig.4. For each observable, a scatter plot is depicted to compare coarse-grained and atomistic results at the smallest force considered, *f* =1 pN. Learning and Testing Sequences are indicated by red circles and green diamonds, respectively, while the black line indicates the bisector of the first and third quadrant. Overall, an excellent agreement is found, as indicated by the localization of the plotted points in the vicinity of the bisector and by the large values attained by the Pearson coefficient for all the considered quantities. In Fig.4a, we plot the crookedness *β*, ^52^ a dimensionless parameter accounting for the global deviation of the helical axis with respect to a straight line, i.e. the presence of a spontaneous curvature. Larger values of *β* correspond to more curved structures, and it was found that A-tracts show the straightest conformations.^52^ In Fig.4b, we analyze the helical diameter. Also in this case, the coarsegrained model faithfully captures the dependence on sequence within a range spanning about 0.2 nm, although a slight overestimation of the atomistic values can be appreciated. In Fig.4c-h we plot the comparison of various helical features, namely h-rise (Fig.4c), h-twist (Fig.4d) and groove geometry (Fig.4e-h). The coarse grained predictions closely follow the sequence-dependence of the atomistic values, although a systematic overestimation of about 0.2 nm is present in the case of the depth of the major groove (Fig.4g). Finally, in Fig.4i we compare the values for the local dihedral SBBS, i.e. the dihedral formed by the sugars and bases within each base pair (inset in Fig.4i). Again, MADna reproduces with high precision the large variability induced by sequence heterogeneity, showing its ability to capture features emerging from the interaction between neighboring base-pairs.

**Figure 4:**
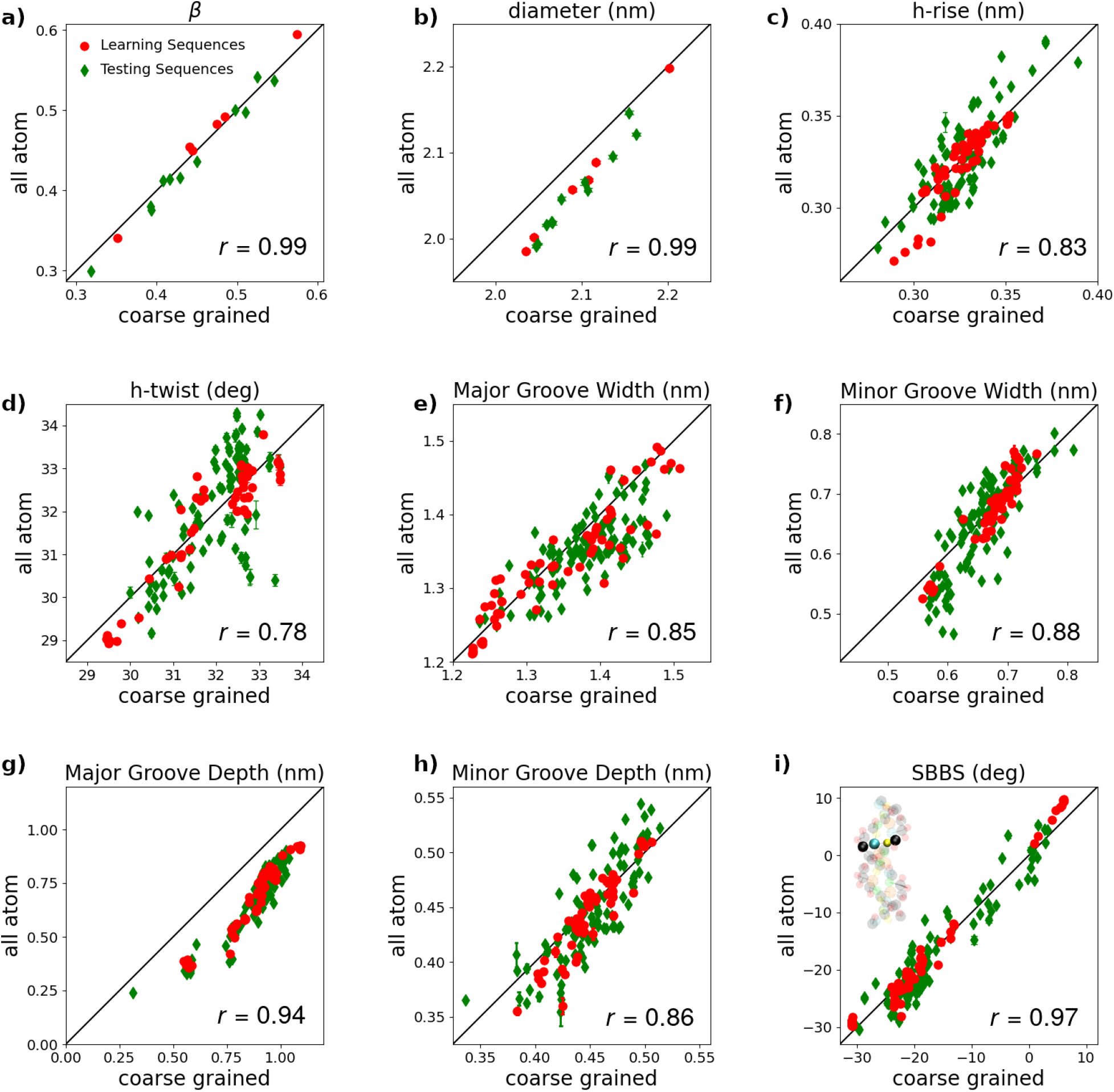
Scatter plot comparing atomistic and coarse-grained results at *f* =1 pN for various structural features: crookedness *β* (panel a); helical diameter (b); helical rise (c); helical twist (d); width of major (e) and minor (f) groove; depth of major (g) and minor (h) groove; SBBS dihedrals (i). Learning and Testing Sequences are denoted by red circles and green diamonds, respectively. Black lines indicate the bisector of the first and third quadrant. For each panel, the Pearson coefficient indicating the linear correlation between the two datasets is reported. The atomistic results were obtained by coarse-graining the trajectories obtained from all-atom simulations and performing the analysis reported in the Methods.

The close agreement between coarse grain and atomistic simulations indicates the high accuracy of MADna in describing the conformational properties of dsDNA. Several striking features are worth mentioning. First, as mentioned above, the Testing Sequences were not employed to build the model, so that in this case the coarse-grained results are pure predictions. Second, none of the quantities reported in Fig.4 were directly used to build the model, thus a quantitative agreement is not trivial also in the case of the Learning Sequences. Third, these observables are related to different scales, encompassing a single base pair (SBBS), a base step (h-twist and h-rise), multiple-step geometry (grooves) and the molecule as a whole (*β* and diameter). Fourth, the selected quantities have a stark dependence on sequence, as shown by their wide range of variability, indicating that the model captures also this intrinsic heterogeneity. Particularly striking is the case of the dihedral SBBS: despite the only two possible choices for the bases (the Watson-Crick pairs AT and CG), this quantity displays a strong variability, ranging from −30 to 10 degrees (Fig.4i). This is evidently an emergent behavior induced by the interaction of these base pairs with their neighbors and is perfectly reproduced by the coarse-grained model.

### MADna reproduces elastic constants obtained from atomistic simulations

Next, we focused on the elastic properties of Learning and Testing Sequences (Fig.5). The effective stretching modulus 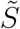 (Fig.5a) and the crookedness elastic constant *k_β_* (Fig.5b) are related to the elastic response of extension and crookedness to the external force, respectively, and are obtained as the slopes of the corresponding observables as a function of *f* (see Methods and Fig.S4 in the Supplementary Information). The torsional modulus *C* (Fig.5c) accounts for the change of the h-twist upon application of an external torque. For the present setup, *C* was computed by analyzing the fluctuations of the h-twist for *f* =1 pN. Finally, the twist-stretch coupling constant g (Fig.5d) quantifies the torsional response to the external force. The negative sign displayed by *g* (Fig.5d) implies that the molecule overwinds when stretched, in agreement with experimental observations.^31,32^ Precise definitions of the elastic constants and their computation are reported in the Methods.

**Figure 5:**
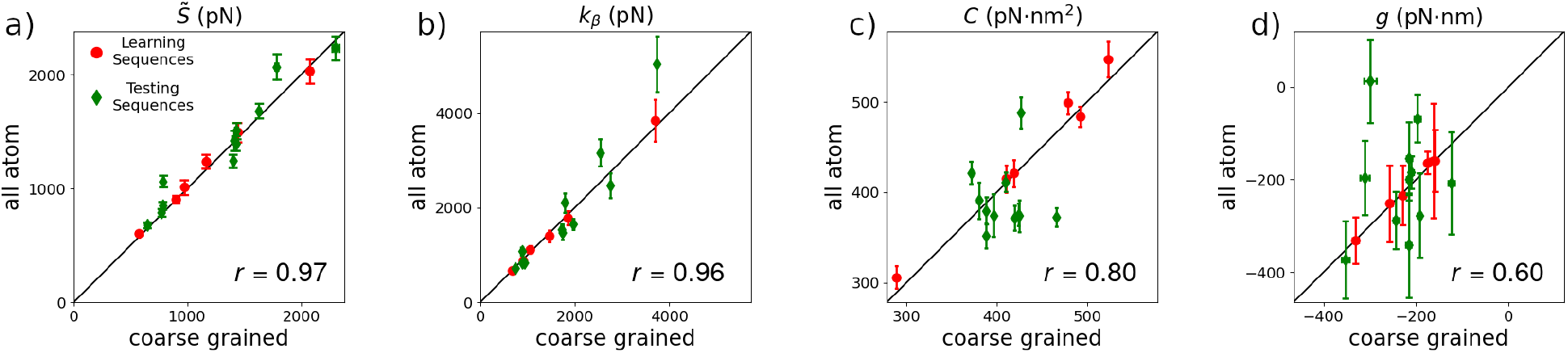
Scatter plot comparing atomistic and coarse-grained results for the effective stretching modulus 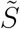 (panel a), the crookedness rigidity *k_β_* (b), the torsion modulus *C* (c) and the twist-stretch coupling constant *g* (d). Learning and Testing Sequences are denoted by red circles and green diamonds, respectively. Black lines indicate the bisector of the first quadrant. For each panel, the Pearson coefficient indicating the linear correlation between the two datasets is reported.

As discussed in the Methods and in Section S2 in the Supplementary Information, the values of 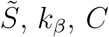 and *g* for the Learning Sequences were used to refine the parameters of the bonded interactions obtained by the Boltzmann Inversion of the atomistic trajectories. Due to the large number of bonded parameters, we opted for a pragmatic approach and adjusted a minimal subset of parameters. Particularly, we empirically observed that a major impact on the elastic constants was obtained by tuning the rigidities of bond 5’-BB-3’, angle 5’-PSB-3’ and dihedral 5’-SPSP-3’. Interestingly, these three bonded interactions have a clear physical meaning (compare Fig.3). Indeed, the bond 5’-BB-3’ accounts for stacking interactions, which are likely to affect the stretching stiffness^52^ as well as the coupling between twist and stretch deformations. Moreover, the range of values encompassed by the angle 5’-PSB-3’ is related to the flexibility of the sugar pucker, which has been shown to play a key role in dsDNA elasticity.^46^ Finally, the 5’-SPSP-3’ dihedral is likely to affect the backbone response and was found to be the quantity most affecting the crookedness rigidity. In this context, it is also worth mentioning that the <u>BB-WC</u> elastic constants for the pairs AT and CG approximately follow the ratio 2:3 (see Table S2 in the Supplementary Information). The <u>BB-WC</u> bonds account for the interactions between Watson-Crick base pairs, hence this ratio nicely reflects the presence of two and three hydrogen bonds for AT and CG, respectively.

As shown in Fig.5, also for the elastic constants a very good agreement is found between coarse-grained and atomistic results (see Tables S5 and S6 in the Supplementary Information for the numerical values), as indicated by the large values of the Pearson coefficient. The only quantity which is somewhat less precisely captured from a quantitative point of view is the twist-stretch coupling *g*, which is however the quantity with the largest indeterminacy. While a good quantitative agreement between the elastic constants obtained in coarse-grained and all-atom simulations is expected for the Learning Sequences (red circles), the good reproduction of the atomistic results also in the case of the Testing Sequences (green diamonds) further confirms that MADna captures the sequence-dependent mechanical properties of dsDNA with precision comparable to atomistic simulations. We note that the values of the elastic constants show wide variations, changing by a factor of two for 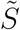 and *C* and reaching four-fold changes for *k_β_* and *g*. It has to be noted that some outliers are present, in particular for sequence A8T the coarse-grained model predicts a twist-stretch coupling markedly different from the atomistic result *g* ≃ 0. Nevertheless, as discussed in Section S6.1 in the Supplementary Information, this disagreement is likely to originate from a possible lack of convergence of the atomistic simulations at large forces for this specific sequence. Based on this observation, we excluded this point for the quantitative evaluation of the agreement between the two datasets by means of the Pearson coefficient reported in Fig.5d.

### Comparison with experiments: sequence-averaged persistence length

The persistence length *l_p_* quantifies the bending rigidity of a polymer and is possibly the most characterized mechanical property of dsDNA.^16–19,21,23,29,37,49^ In its classical formulation from polymer theory, *l_p_* corresponds to the length of the fragment to be considered in order to observe significant thermally-induced bending effects. ^90^ More technically, *l_p_* is defined as the decay length of the thermally-averaged tangent-vector correlation function *c_p_*, that is *c_p_* = exp(–*s*/*l_p_*), where *s* is the contour length of the polymer fragment under consideration.^90^ This definition relies on the assumption that the minimum-energy conformation for the polymer is that of a straight rod. In the case of dsDNA, according to sequence a spontaneous curvature may be present. In order to account separately for intrinsic bending and thermal fluctuations, it is customary to distinguish the static (*l_s_*) and dynamic (*l_d_*) contributions to the persistence length, which characterize the decay length of the suitably-defined correlation functions *c_s_* = exp(−*s/l_s_*) (computed from the average structure) and *c_d_* = exp(−*s*/*l_d_*).^56^ The three lengths are approximately related by a harmonic sum, 1/*l_p_* ≃ 1/*l_s_* + 1/*l_d_*.^87^

In order to compute the persistence length and its contributions, we performed simulations for 20 random sequences of length 100 base pairs. The average structures computed by the molecular builder (see Methods) and the equilibrated trajectories were then employed to compute the correlation functions *c_p_, c_s_* and *c_d_* according to Equations (12), (13) and (14), respectively. The results are reported in Fig.S6 in the Supplementary Information for various definitions of the tangent vector. Here, we focus on the results of one of such definitions, according to which the tangent vectors are obtained by joining the geometrical centers of sugars of base pairs separated by ten steps (see Methods). Fitting the correlation function *c_p_* with an exponential decay resulted in computed values of the persistence length *l_p_* ranging between 46 nm and 64 nm, with an average equal to *l_p_* = 56 ± 1 nm. This is in good agreement with experimental values on random sequences and standard ionic conditions (45-55 nm^16–19,21,23,29,37,49^), particularly since *l_p_* was not employed in the parameterization of the model. Other coarse-grained models give predictions for *l_p_* within the experimental range,^60,62,65,68,70^ although in most of these works the persistence length of double-^60,68,70^ or single-stranded^65^ DNA was employed as a target quantity in the construction of the force field. In the present case, the slight overestimation of *l_p_* is in line with previous results of atomistic simulations, where an average *l_p_* = 57 ± 3 was found.^49^

As expected, ^56^ for each sequence the static correlation function *c_s_* is found to decrease more slowly than *c_p_* (Fig.S6 in the Supplementary Information), since it lacks the disordering action of thermal fluctuations. The corresponding static persistence length *l_s_* was computed by fitting *c_s_* via an exponential decay, in analogy to *c_p_*. The obtained values vary widely, ranging from 101 nm to 786 nm, with an average equal to *l_s_* = 325 ± 47 nm. This is somewhat smaller than previous estimations (*l_s_* = 576 ± 191 nm^49^), although we note that this value varies wildly by considering different sequences and, even for the same sequences, by employing different definitions (see below). As discussed in Ref.,^49^ its large heterogeneity may rationalize the markedly-different results reported in experiments, where values of *l_s_* as different as 130 nm and >1000 nm have been estimated.^16,91^

Fitting *c_d_* via an exponential decay yields values for the dynamic persistence length *l_d_* ranging between 62 nm and 116 nm, with an average equal to *l_d_* = 75 ± 3 nm. This prediction is in agreement with the experimental value *l_d_* = 82 ± 15 nm obtained in Ref.,^16^ although departing from the measurements from Ref.,^91^ for which *l_d_* = 50 ± 1 nm. The difference between the two experimental values might be ascribed to the different techniques employed, for which future simulations with the present model may shed some light. When compared to atomistic predictions (*l_d_* = 64.7± 1.4 nm^49^), the value found here is a bit larger. This is coherent with the smaller value found above for the static persistence *l_s_*. Indeed, from the Trifonov relation 1/*l_p_* = 1/*l_S_* + 1/*l_d_*, in order to obtain similar values for *l_p_* the difference observed for *l_s_* needs to be compensated by an opposite behavior for *l_d_*.

As a final consideration, it has to be noted that the estimation of the persistence length may vary according to the definition of the tangent vectors. ^56^ Alternative choices are considered in Section S6 in the Supplementary Information, yielding average values in the ranges *l_p_* = 56 – 63 nm, *l_s_* = 326 – 633 nm and *l_d_* = 75 – 77 nm. This indicates the robustness of the values of *l_p_* and *l_d_* with respect to the definition of the tangent vectors. The average value of ls appears to be more sensitive to the definition, although the observed variation is relatively small when compared to the intrinsic large variability characterizing this quantity.

### Comparison with experiments: sequence-averaged elastic constants

A further set of simulations was devoted to determine the sequence-averaged elastic constants. These quantities were already estimated for the Learning and Testing Sequences (Fig.5), but their quantitative determination is likely to depend on the details of the microscopic definitions, particularly for the h-twist. In order to enable a quantitative comparison with experimental values, we designed a simulation framework which avoids relying on such definitions and which is more akin to single-molecule experimental setups.

As shown schematically in Fig.6a, in this set of simulations a dsDNA molecule is subjected to a constant force *f* and torque *τ* acting along the z axis. This enables defining unequivocally the twist angle *θ* starting from the projection of the base pairs onto the xy plane, in analogy with single-molecule experiments based on rotor beads. ^31^ The linear response of the extension *L* as a function of *f* provides access to the effective stretch modulus 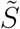. Analogously, the torsion modulus *C* can be computed from the dependence of *θ* on *τ*. Finally, the twist-stretch coupling *g* can be obtained by looking at the cross-dependence, i.e. the dependence of *L* on *τ* or the response of *θ* to changes in *f*. The quantitative details can be found in the Methods.

**Figure 6:**
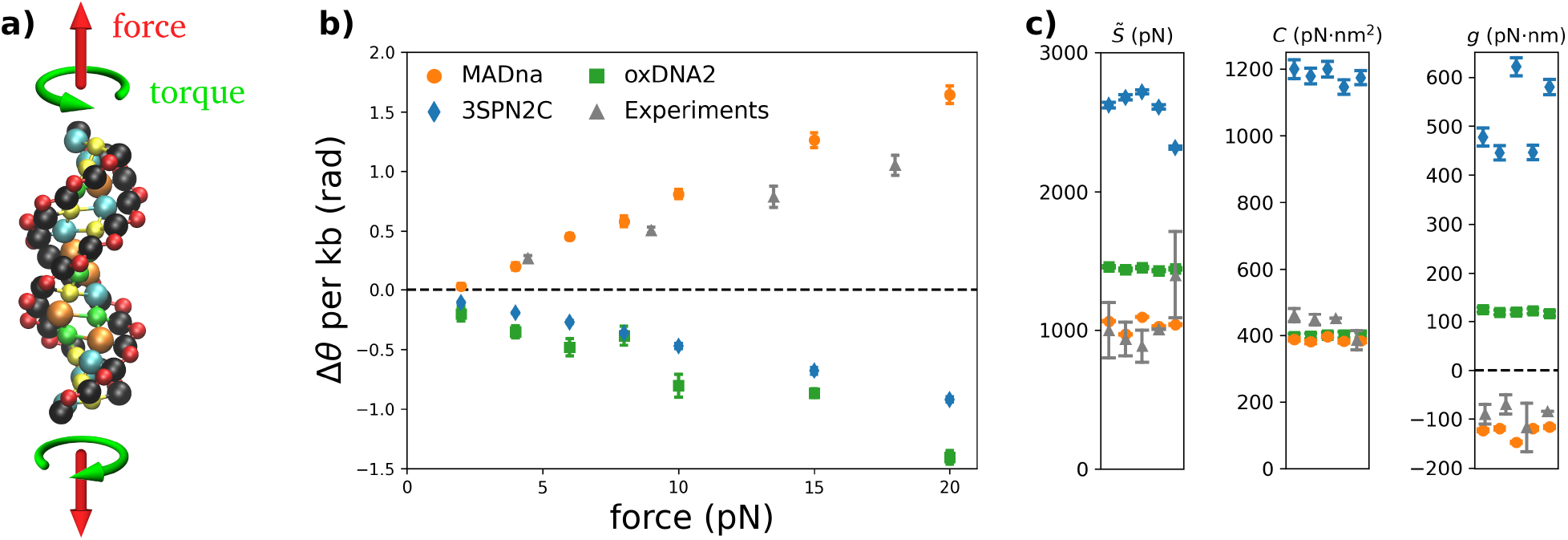
a) Schematic description of the simulation setup for the stretch-torsion simulations, prescribing a constant force and torque applied along a fixed direction. b) Twist response to the external force in the absence of imposed torque for MADna (orange circles), oxDNA2 (green squares) 3SPN2C (blue diamonds) and rotor-bead experiments^31^ (grey triangles). The black dashed line indicates Δ*θ* = 0, therefore overwinding and unwinding responses are characterized by points lying above and below the line, respectively. The simulation data correspond to the average of the five sequences considered. c) Effective stretching modulus 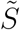, torsion modulus *C* and twist-stretch coupling constant *g* obtained by the three models for the five sequences reported in Section S3.3 in the Supplementary Information. Symbols are the same as in panel b. The grey triangles correspond to experimental measures obtained for unrelated sequences in Refs.^17–20^ for 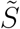, Refs.^20,27,31,92^ for *C* and Refs.^20,31,32,93^ for *g*. In the case of *g*, the dashed black line denotes *g* = 0, thus separating two regimes characterized by qualitatively-different twist-stretch coupling.

We performed simulations for five randomly-generated sequences of length equal to forty base pairs at several values of *f* and *τ* (see Methods for full details and Section S3.3 in the Supplementary Information for the sequences, which are named ST1,…,ST5). With this choice of sequence length, the molecules are long enough so that several turns of the double helix are present, but short enough to neglect the effect of bending. For comparison, we also run simulations for the same set of sequences and parameters by employing the two most-widely used coarse-grained models from literature, namely oxDNA2 ^80,81^ and the sequence-dependent model 3SPN2C. ^82^ From a qualitative perspective, a marked difference between MADna and the other two models becomes evident when analyzing the change in twist Δ*θ* as a function of the force (Fig.6b). Indeed, the present model prescribes that dsDNA overwinds when stretched (Δ*θ* > 0), which is in agreement with experimental observations.^31,32^ In contrast, neither oxDNA2 nor 3SPN2C capture this feature, predicting unwinding upon pulling (in the case of oxDNA this fact was already observed in the original publication^68^). To our knowledge, no other coarse-grained model available in literature has been shown to predict this unintuitive behavior of dsDNA.

MADna shows better agreement with experiments also from a quantitative standpoint. As shown in Fig.6c and Table 1, it predicts values for 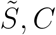 and *g* in good quantitative agreement with the experiments. As for the other models, oxDNA2 shows a similar performance for *C*, while it tends to overestimate 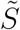. Coherently with the results shown in Fig.6b, the wrong sign for *g* is found for both oxDNA2 and 3SPN2C, with the latter showing in general a significant overestimation of the elastic constants. The results reported in Fig.6c and Table 1 correspond to forces larger than 10 pN, which was chosen as a reasonable threshold to avoid bending effects. Nevertheless, performing the analysis with the full set of forces resulted in a small change in 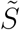 (roughly 10%) and in virtually no change in *C* and *g*. It is also worth mentioning that the range considered for the torque (0 – 30 pN·nm) corresponds to supercoiling densities below the threshold value *σ* ≃ 0.05 usually associated to torque-induced denaturation, ^22^ hence making the results obtained for MADna relevant for real dsDNA molecules within the whole range of applied torques. Particularly, we computed the supercoiling density as *σ* = Δ*θ*/*θ*_0_, since the lack of bending results in absence of relevant writhe. For the largest torque applied (*τ* = 30 pN·nm), we found *σ* ≃ 0.046 for MADna, *σ* ≃ 0.043 for oxDNA2 and *σ* ≃ 0.014 for 3SPN2C, which is lower than for the other two models due to the larger value of the twist modulus *C*.

**Table 1:** Elastic constants obtained by the various models and their comparison with experiments (cfr Fig.6c). Experimental values are obtained as averages of the results reported in Refs.^17–20^ for 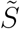, Refs.^20,27,31,92^ for *C* and Refs.^20,31,32,93^ for *g*, while the corresponding indeterminacies are computed as standard error of the mean.

| | $\tilde{S}$ (pN) | $C$ (pN·nm<sup>2</sup>) | $g$ (pn·nm) |
| --- | --- | --- | --- |
| Experiment | $1045 \pm 92$ | $436 \pm 17$ | $-90 \pm 10$ |
| MADna | $1038 \pm 21$ | $386 \pm 3$ | $-125 \pm 6$ |
| oxDNA2 | $1448 \pm 5$ | $399 \pm 1$ | $+120 \pm 1$ |
| 3SPN2C | $2589 \pm 71$ | $1180 \pm 10$ | $+514 \pm 36$ |

Comparing the results obtained for MADna in Fig.5 and Fig.6 indicates a marked discrepancy between the values found for *g*, which change by as much as a factor of two. This striking disagreement appears to be too large to be simply ascribed to the different sequences considered or the different definitions employed for the twist angle. In this regard, based on atomistic simulations it has been suggested that the stretching modulus depends on the size of the fragment under consideration. ^94^ Since in the present case we are comparing sequences of size twenty and fourty base pairs, it might be that a similar effect is present for *g*. In order to check this hypothesis, we performed another set of simulations following the same protocol as in the present section, but considering sequences made of twenty base pairs, which are listed as ST1-short,…,ST5-short in Section S3.3 in the Supplementary Information. The resulting elastic constants are reported in Fig.S7 in the Supplementary Information. Particularly, we found that for the short sequences *g* = −258 ± 16 pN·nm, which is indeed in line with the results reported in Fig.5. This confirms the presence of size effects in MADna for the prediction of elastic constants. To our knowledge, this is the first instance in which a length dependence for *g* has been predicted. This feature might explain the systematic overestimation of *g* in atomistic simulations that are restricted to short sequences. ^46^

Another interesting effect observed in experiments is the coupling between twist and bending, which is relevant at pulling forces up to a few pN. ^33,34^ As a consequence of this coupling, the twist of the DNA molecule appears to be softer, i.e. the effective twist modulus is *C*_eff_ < *C*. In order to study this feature, we considered a set of simulations involving DNA molecules of 150 base pairs, which is comparable to the persistence length, thus enabling the presence of bending fluctuations. The simulation protocol was similar to the case just analyzed, although here no torque was applied. We considered three independent sequences which are reported in Section S3.4 in the Supplementary Information. For each sequence, simulations were performed for MADna, oxDNA2 and 3SPN2C at pulling forces ranging between 0.05 pN and 2.5 pN. Further simulation details can be found in the Methods. In Fig.7 we report the results obtained and compare them with the experimental values from Refs.^33,34^ In line with the trend observed in Fig.6, MADna and oxDNA2 follow quite closely the experimental curve, while 3SPN2C systematically overestimates the value of *C*_eff_.

**Figure 7:**
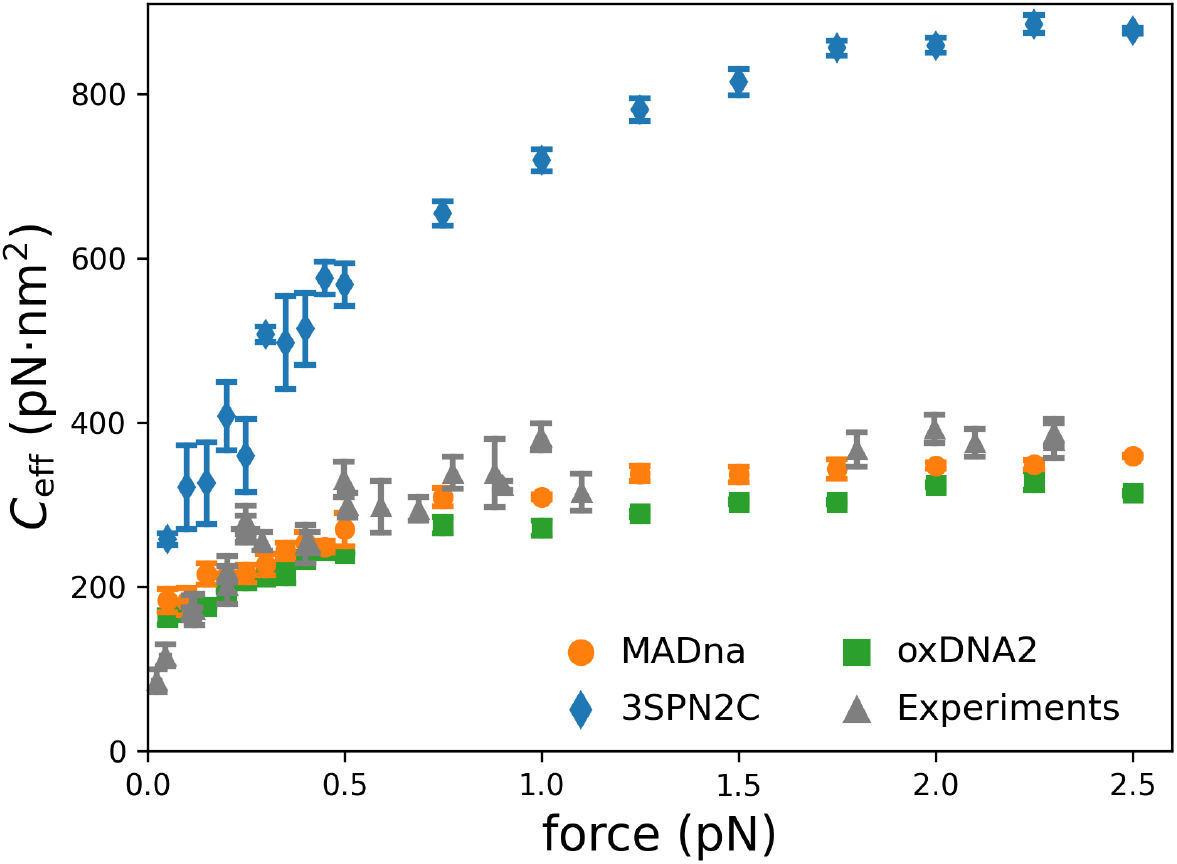
Effective torsion modulus *C*_eff_ as a function of force for MADna (orange circles), oxDNA2 (green squares), 3SPN2C (blue diamonds) and experiments (grey triangles). Experimental data were extracted from Refs.^33,34^

### Comparison with experiments: sequence-dependent conformation and elasticity

Having characterized and benchmarked the sequence-averaged mechanical properties of MADna, we now turn to sequence-dependent features. In this regard, we considered the sequences studied experimentally in Ref., ^28^ where the authors characterized the sequence-dependent persistence length and helical repeat by means of cyclization experiments.

We performed simulations for DNA molecules of 100 base pairs obtained by taking the central parts of 14 different experimental sequences as listed in Section S3.2 in the Supplementary Information. For each simulated sequence, we computed the helical pitch by dividing the cumulated helical twist by 2π, while for the persistence length we employed the same approach as above. In Fig.8 we report the comparison between experiments and MADna predictions. As the plots show, there is a high correlation between the two datasets, thus indicating that MADna satisfactorily captures the sequence dependence of two key features of DNA conformation (helical pitch) and elasticity (persistence length). From a quantitative perspective, we see from Fig.8a that the simulated values for the pitch are larger than the experimental ones. Nonetheless, a quantitative comparison has to be performed with care. Indeed, the experimental values were obtained indirectly by analyzing cyclization data by means of a wormlike chain with twist. ^28^ Although the relative differences observed in the experimental results for the different sequences are robust, the values depend on the validity of this model down to the scale of single steps and on the value assigned to the twist modulus when fitting the data. Moreover, it is likely that the details of the definition of the h-twist in the simulations affect the quantitative determination of the helical pitch. To our knowledge, this is the first instance in which these structural data have been compared to predictions from simulations. As for the persistence length, the strong correlation found has to be mitigated by the slight quantitative overestimation of lp in simulations, in line with the results reported above for the sequence-averaged persistence length.

**Figure 8:**
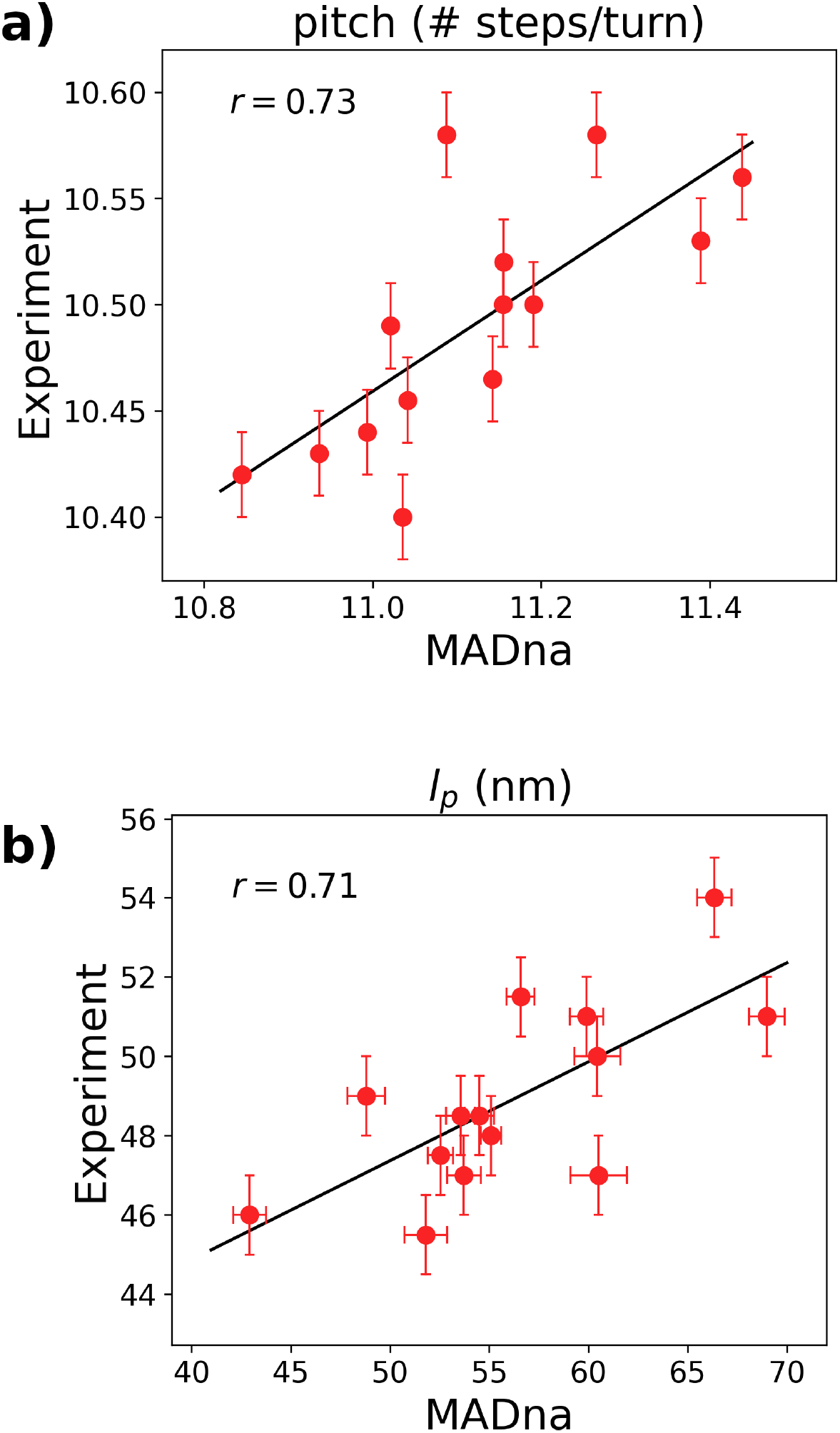
Comparison between experimental values and MADna predictions for the sequence dependence of the helical pitch (a) and the persistence length *l_p_* (b). The lines correspond to the linear fits of the scatter plots and are included as a guide to the eye. The value of the Pearson coefficient is reported in each plot.

For comparison, we also performed simulations for oxDNA2 and 3SPN2C, for which the results are reported in Fig.S8 in the Supplementary Information. We found that oxDNA2 predictions do not correlate with experiments for neither the helical pitch nor the persistence length. This is expected, since in this model the sequence dependence is implemented by tuning the base-pairing and stacking interactions to account for thermodynamic data, but not for the elasticity of DNA. In the case of 3SPN2C, we found a weak correlation for the helical pitch but a strong correlation for the persistence length. This high correlation was also expected, since the persistence-length data from Ref. ^28^ were used to parameterize the model. ^82^ For completeness, we mention that also CGDNA has been used to reproduce these experimental persistence-length data, for which a similar correlation as for MADna was found (*r* = 0.73).^56^

As a further test, we performed stretching simulations for phased A-tracts, i.e. dsDNA molecules obtained by alternating fragments of consecutive adenines and random sequences, with each fragment having a length of 5-10 base pairs. It was experimentally found that the stretching modulus for such molecules is roughly 50% larger than for random sequences.^21^ We run some stretching simulations for five phased A-tracts of 40 base pairs, whose sequences were extracted as fragments of the experimental ones^21^ and are reported as A-tract-1,…, A-tract-5 in Section S3.3 in the Supplementary Information. Simulation protocols and analysis were the same as in the case of random sequences studied above. The results from the three models were 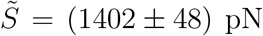 for MADna, 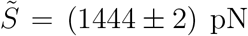 for oxDNA2 and 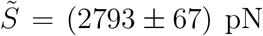 for 3SPN2C. By comparing these values with the results reported in Table 1, we thus conclude that MADna predicts an increase of roughly 35%, oxDNA2 does not predict any increase and 3SPN2C predicts a 8% increase. As expected, the absence of elasticity-oriented sequence dependence for oxDNA2 prevents it from capturing this feature. The prediction of MADna is the one most closely resembling the significant increase observed experimentally, although still unnderestimating its extent.

### Strengths and limits

The comparison between MADna, oxDNA2, 3SPN2C and experiments can be summarized as follows. MADna performs in general better at capturing both sequence-averaged and sequence-dependent conformational and elastic features. As for the other two models, oxDNA2 reproduces well the sequence-averaged properties, but it cannot account for sequence dependence. In contrast, 3SPN2C captures well the sequence dependence of elasticity in the case of the persistence length, while its accuracy for the change in stretch modulus or conformation is more limited. Moreover, it tends in general to overestimate the magnitude of the elastic constants.

The accuracy of MADna in addressing the sequence dependence of conformational and elastic properties of dsDNA appoints it as the ideal choice to interpret the outcome of singlemolecule experiments, as well as to study the conformational changes induced by mechanic stress, which is relevant for many systems in vivo. Moreover, it can be employed as a virtual laboratory to test analytical theories on DNA elasticity. ^95,96^ Nonetheless, at the present stage MADna cannot account for breaking events such as the formation of kinks or local melting. Hence, it is important to assess the relevance of such mechanisms for the system under study before drawing conclusions based on simulations performed with the present model. We are currently working to surpass such limitations, in order to further extend the palette of possible systems which can be analyzed through the lens of MADna.

## Conclusion

We have introduced MADna, a novel coarse-grained model for the simulation of doublestranded DNA. MADna captures the sequence dependence of conformational and elastic properties with accuracy comparable to that of atomistic simulations. Key conformational features which closely follow atomistic results include the main helical parameters, the groove geometry, the diameter of the double helix and the spontaneous curvature as quantified by the crookedness. The model also predicts sequence-averaged and sequence-dependent elasticity and conformation in agreement with experimental results for a wide set of features, namely the stretching and torsion moduli, the counterintuitive negative twist-stretch coupling, the twist-bend coupling, the persistence length and the helical pitch. At the present stage, the model does not account for breaking events, which are being addressed in ongoing work. Implementation of MADna in Molecular Dynamics softwares is straightforward due to the common potentials employed.

Due to both its accuracy and its simplicity of use, we believe that MADna provides a significant addition to the toolbox of coarse-grained simulations, and will enable precise theoretical studies of a wide set of large-scale DNA phenomena.

## Acknowledgements

We are indebted to Alberto Marín-González and Alessandro Barducci for insightful discussions. We thank the financial support from the Spanish MINECO project MAT2017-83273-R. We acknowledge support from the Ministerio de Ciencia e Innovación (MICINN) through the project PID2020-115864RB-I00 and the “María de Maeztu” Programme for Units of Excellence in R&D (grant No. CEX2018-000805-M). The project that gave rise to these results received the support of a fellowship from “la Caixa” Foundation (ID 100010434) and from the European Union’s Horizon research and innovation programme under the Marie Skłodowska-Curie grant agreement No 847648. The fellowship code is LCF/BQ/PI20/11760019.

## References

(1) Rohs, R.; Jin, X.; West, S. M.; Joshi, R.; Honig, B.; Mann, R. S. Origins of specificity in protein-DNA recognition. Annual review of biochemistry 2010, 79, 233–269.

(2) Rohs, R.; West, S. M.; Sosinsky, A.; Liu, P.; Mann, R. S.; Honig, B. The role of DNA shape in protein–DNA recognition. Nature 2009, 461, 1248–1253.

(3) Segal, E.; Widom, J. Poly (dA: dT) tracts: major determinants of nucleosome organization. Current opinion in structural biology 2009, 19, 65–71.

(4) Haran, T. E.; Mohanty, U. The unique structure of A-tracts and intrinsic DNA bending. Quarterly reviews of biophysics 2009, 42, 41–81.

(5) Seol, Y.; Hardin, A. H.; Strub, M.-P.; Charvin, G.; Neuman, K. C. Comparison of DNA decatenation by Escherichia coli topoisomerase IV and topoisomerase III: implications for non-equilibrium topology simplification. Nucleic acids research 2013, 41, 4640–4649.

(6) Dillon, S. C.; Dorman, C. J. Bacterial nucleoid-associated proteins, nucleoid structure and gene expression. Nature Reviews Microbiology 2010, 8, 185–195.

(7) Saha, A.; Wittmeyer, J.; Cairns, B. R. Chromatin remodelling: the industrial revolution of DNA around histones. Nature reviews Molecular cell biology 2006, 7, 437–447.

(8) Lee, J. Y.; Terakawa, T.; Qi, Z.; Steinfeld, J. B.; Redding, S.; Kwon, Y.; Gaines, W. A.; Zhao, W.; Sung, P.; Greene, E. C. Base triplet stepping by the Rad51/RecA family of recombinases. Science 20 1 5, 349, 977–981.

(9) Ong, M. S.; Richmond, T. J.; Davey, C. A. DNA stretching and extreme kinking in the nucleosome core. Journal of molecular biology 2007, 368, 1067–1074.

(10) Dikic, J.; Seidel, R. Anticooperative binding governs the mechanics of ethidium-complexed DNA. Biophysical journal 2019, 116, 1394–1405.

(11) Chhabra, H.; Mishra, G.; Cao, Y.; Prešern, D.; Skoruppa, E.; Tortora, M. M.; Doye, J. P. Computing the Elastic Mechanical Properties of Rodlike DNA Nanostructures. Journal of Chemical Theory and Computation 2020, 16, 7748–7763.

(12) Mathew-Fenn, R. S.; Das, R.; Harbury, P. A. Remeasuring the double helix. Science 2008, 322, 446–449.

(13) Shi, X.; Herschlag, D.; Harbury, P. A. Structural ensemble and microscopic elasticity of freely diffusing DNA by direct measurement of fluctuations. Proceedings of the National Academy of Sciences 2013, 110, E1444–E1451.

(14) Ido, S.; Kimura, K.; Oyabu, N.; Kobayashi, K.; Tsukada, M.; Matsushige, K.; Yamada, H. Beyond the helix pitch: direct visualization of native DNA in aqueous solution. ACS nano 2013, 7, 1817–1822.

(15) Kuchuk, K.; Katrivas, L.; Kotlyar, A.; Sivan, U. Sequence-dependent deviations of constrained DNA from canonical B-form. Nano letters 2019, 19, 6600–6603.

(16) Bednar, J.; Furrer, P.; Katritch, V.; Stasiak, A.; Dubochet, J.; Stasiak, A. Determination of DNA persistence length by cryo-electron microscopy. Separation of the static and dynamic contributions to the apparent persistence length of DNA. Journal of molecular biology 1995, 254, 579–594.

(17) Baumann, C. G.; Smith, S. B.; Bloomfield, V. A.; Bustamante, C. Ionic effects on the elasticity of single DNA molecules. Proceedings of the National Academy of Sciences 1997, 94, 6185–6190.

(18) Wenner, J. R.; Williams, M. C.; Rouzina, I.; Bloomfield, V. A. Salt dependence of the elasticity and overstretching transition of single DNA molecules. Biophysical journal 2002, 82, 3160–3169.

(19) Herrero-Galán, E.; Fuentes-Perez, M. E.; Carrasco, C.; Valpuesta, J. M.; Carrascosa, J. L.; Moreno-Herrero, F.; Arias-González, J. R. Mechanical identities of RNA and DNA double helices unveiled at the single-molecule level. Journal of the American Chemical Society 2013, 135, 122–131.

(20) Lipfert, J.; Skinner, G. M.; Keegstra, J. M.; Hensgens, T.; Jager, T.; Dulin, D.; Kober, M.; Yu, Z.; Donkers, S. P.; Chou, F.-C., et al. Double-stranded RNA under force and torque: similarities to and striking differences from double-stranded DNA. Proceedings of the National Academy of Sciences 2014, 111, 15408–15413.

(21) Marin-González, A.; Pastrana, C. L.; Bocanegra, R.; Martín-Gonzalez, A.; Vilhena, J.; Píerez, R.; Ibarra, B.; Aicart-Ramos, C.; Moreno-Herrero, F. Understanding the paradoxical mechanical response of in-phase A-tracts at different force regimes. Nucleic acids research 2020, 48, 5024–5036.

(22) Bryant, Z.; Stone, M. D.; Gore, J.; Smith, S. B.; Cozzarelli, N. R.; Bustamante, C. Structural transitions and elasticity from torque measurements on DNA. Nature 2003, 424, 338–341.

(23) Lipfert, J.; Kerssemakers, J. W.; Jager, T.; Dekker, N. H. Magnetic torque tweezers: measuring torsional stiffness in DNA and RecA-DNA filaments. Nature methods 2010, 7, 977–980.

(24) Lipfert, J.; Wiggin, M.; Kerssemakers, J. W.; Pedaci, F.; Dekker, N. H. Freely orbiting magnetic tweezers to directly monitor changes in the twist of nucleic acids. Nature communications 2011, 2, 1–10.

(25) Kriegel, F.; Ermann, N.; Forbes, R.; Dulin, D.; Dekker, N. H.; Lipfert, J. Probing the salt dependence of the torsional stiffness of DNA by multiplexed magnetic torque tweezers. Nucleic acids research 2017, 45, 5920–5929.

(26) Vanderlinden, W.; Kolbeck, P. J.; Kriegel, F.; Walker, P. U.; Lipfert, J. A benchmark data set for the mechanical properties of double-stranded DNA and RNA under torsional constraint. Data in brief 2020, 30, 105404.

(27) Mosconi, F.; Allemand, J. F.; Bensimon, D.; Croquette, V. Measurement of the torque on a single stretched and twisted DNA using magnetic tweezers. Physical review letters 2009, 102, 078301.

(28) Geggier, S.; Vologodskii, A. Sequence dependence of DNA bending rigidity. Proceedings of the National Academy of Sciences 2010, 107, 15421–15426.

(29) Guilbaud, S.; Salomé, L.; Destainville, N.; Manghi, M.; Tardin, C. Dependence of DNA persistence length on ionic strength and ion type. Physical review letters 2019, 122, 028102.

(30) Heenan, P. R.; Perkins, T. T. Imaging DNA equilibrated onto mica in liquid using biochemically relevant deposition conditions. ACS nano 2019, 13, 4220–4229.

(31) Gore, J.; Bryant, Z.; Nöllmann, M.; Le, M. U.; Cozzarelli, N. R.; Bustamante, C. DNA overwinds when stretched. Nature 2006, 442, 836–839.

(32) Gross, P.; Laurens, N.; Oddershede, L. B.; Bockelmann, U.; Peterman, E. J.; Wuite, G. J. Quantifying how DNA stretches, melts and changes twist under tension. Nature Physics 2011, 7, 731–736.

(33) Nomidis, S. K.; Kriegel, F.; Vanderlinden, W.; Lipfert, J.; Carlon, E. Twist-bend coupling and the torsional response of double-stranded DNA. Physical review letters 2017, 118, 217801.

(34) Gao, X.; Hong, Y.; Ye, F.; Inman, J. T.; Wang, M. D. Torsional stiffness of extended and plectonemic DNA. Physical Review Letters 2021, 127, 028101.

(35) Lionnet, T.; Joubaud, S.; Lavery, R.; Bensimon, D.; Croquette, V. Wringing out DNA. Physical Review Letters 2006, 96, 178102.

(36) Wiggins, P. A.; Van Der Heijden, T.; Moreno-Herrero, F.; Spakowitz, A.; Phillips, R.; Widom, J.; Dekker, C.; Nelson, P. C. High flexibility of DNA on short length scales probed by atomic force microscopy. Nature nanotechnology 2006, 1, 137–141.

(37) Mazur, A. K.; Maaloum, M. Atomic force microscopy study of DNA flexibility on short length scales: smooth bending versus kinking. Nucleic acids research 2014, 42, 14006–14012.

(38) Vafabakhsh, R.; Ha, T. Extreme bendability of DNA less than 100 base pairs long revealed by single-molecule cyclization. Science 2012, 337, 1097–1101.

(39) Le, T. T.; Kim, H. D. Probing the elastic limit of DNA bending. Nucleic acids research 2014, 42, 10786–10794.

(40) Jeong, J.; Kim, H. D. Determinants of cyclization–decyclization kinetics of short DNA with sticky ends. Nucleic acids research 2020, 48, 5147–5156.

(41) Basu, A.; Bobrovnikov, D. G.; Qureshi, Z.; Kayikcioglu, T.; Ngo, T. T.; Ranjan, A.; Eustermann, S.; Cieza, B.; Morgan, M. T.; Hejna, M.; Rube, H. T.; Hopfner, K.-P.; Wolberger, C.; Song, J. S.; Ha, T. Measuring DNA mechanics on the genome scale. Nature 2021, 589, 462–467.

(42) Becker, N. B.; Everaers, R. Comment on “Remeasuring the double helix”. Science 2009, 325, 538–538.

(43) Mathew-Fenn, R. S.; Das, R.; Fenn, T. D.; Schneiders, M.; Harbury, P. A. Response to comment on “Remeasuring the double helix”. Science 2009, 325, 538–538.

(44) Harrison, R. M.; Romano, F.; Ouldridge, T. E.; Louis, A. A.; Doye, J. P. Identifying physical causes of apparent enhanced cyclization of short DNA molecules with a coarsegrained model. Journal of chemical theory and computation 2019, 15, 4660–4672.

(45) Pasi, M.; Lavery, R. Structure and dynamics of DNA loops on nucleosomes studied with atomistic, microsecond-scale molecular dynamics. Nucleic acids research 2016, 44, 5450–5456.

(46) Marin-Gonzalez, A.; Vilhena, J.; Perez, R.; Moreno-Herrero, F. Understanding the mechanical response of double-stranded DNA and RNA under constant stretching forces using all-atom molecular dynamics. Proceedings of the National Academy of Sciences 2017, 114, 7049–7054.

(47) Bao, L.; Zhang, X.; Shi, Y.-Z.; Wu, Y.-Y.; Tan, Z.-J. Understanding the relative flexibility of RNA and DNA duplexes: stretching and twist-stretch coupling. Biophysical journal 2017, 112, 1094–1104.

(48) Kriegel, F.; Matek, C.; Dršata, T.; Kulenkampff, K.; Tschirpke, S.; Zacharias, M.; Lankaš, F.; Lipfert, J. The temperature dependence of the helical twist of DNA. Nucleic acids research 2018, 46, 7998–8009.

(49) Velasco-Berrelleza, V.; Burman, M.; Shepherd, J. W.; Leake, M. C.; Golestanian, R.; Noy, A. SerraNA: a program to determine nucleic acids elasticity from simulation data. Physical Chemistry Chemical Physics 2020, 22, 19254–19266.

(50) Pasi, M.; Maddocks, J. H.; Beveridge, D.; Bishop, T. C.; Case, D. A.; Cheatham III, T.; Dans, P. D.; Jayaram, B.; Lankas, F.; Laughton, C., et al. *μ*ABC: a systematic microsecond molecular dynamics study of tetranucleotide sequence effects in B-DNA. Nucleic acids research 2014, 42, 12272–12283.

(51) Zgarbová, M.; Jurecka, P.; Lankas, F.; Cheatham III, T. E.; Sponer, J.; Otyepka, M. Influence of BII backbone substates on DNA twist: a unified view and comparison of simulation and experiment for all 136 distinct tetranucleotide sequences. Journal of chemical information and modeling 2017, 57, 275–287.

(52) Marin-Gonzalez, A.; Vilhena, J.; Moreno-Herrero, F.; Perez, R. DNA crookedness regulates DNA mechanical properties at short length scales. Physical review letters 2019, 122, 048102.

(53) Shepherd, J. W.; Greenall, R. J.; Probert, M. I. J.; Noy, A.; Leake, M. C. The emergence of sequence-dependent structural motifs in stretched, torsionally constrained DNA. Nucleic acids research 2020, 48, 1748–1763.

(54) Petkevičiūtė, D.; Pasi, M.; Gonzalez, O.; Maddocks, J. cgDNA: a software package for the prediction of sequence-dependent coarse-grain free energies of B-form DNA. Nucleic acids research 2014, 42, e153–e153.

(55) Walther, J.; Dans, P. D.; Balaceanu, A.; Hospital, A.; Bayarri, G.; Orozco, M. A multimodal coarse grained model of DNA flexibility mappable to the atomistic level. Nucleic acids research 2020, 48, e29–e29.

(56) Mitchell, J. S.; Glowacki, J.; Grandchamp, A. E.; Manning, R. S.; Maddocks, J. H. Sequence-dependent persistence lengths of DNA. Journal of chemical theory and computation 2017, 13, 1539–1555.

(57) Knotts IV, T. A.; Rathore, N.; Schwartz, D. C.; De Pablo, J. J. A coarse grain model for DNA. The Journal of chemical physics 2007, 126, 02B611.

(58) Ouldridge, T. E.; Louis, A. A.; Doye, J. P. DNA nanotweezers studied with a coarsegrained model of DNA. Physical Review Letters 20 1 0, 104, 178101.

(59) Morriss-Andrews, A.; Rottler, J.; Plotkin, S. S. A systematically coarse-grained model for DNA and its predictions for persistence length, stacking, twist, and chirality. The Journal of chemical physics 2010, 132, 01B611.

(60) Savelyev, A.; Papoian, G. A. Chemically accurate coarse graining of double-stranded DNA. Proceedings of the National Academy of Sciences 2010, 107, 20340–20345.

(61) Cragnolini, T.; Derreumaux, P.; Pasquali, S. Coarse-grained simulations of RNA and DNA duplexes. The Journal of Physical Chemistry B 2013, 117, 8047–8060.

(62) Korolev, N.; Luo, D.; Lyubartsev, A. P.; Nordenskiöld, L. A coarse-grained DNA model parameterized from atomistic simulations by inverse Monte Carlo. Polymers 2014, 6, 1655–1675.

(63) Maciejczyk, M.; Spasic, A.; Liwo, A.; Scheraga, H. A. DNA duplex formation with a coarse-grained model. Journal of chemical theory and computation 2014, 10, 5020–5035.

(64) Uusitalo, J. J.; Ingólfsson, H. I.; Akhshi, P.; Tieleman, D. P.; Marrink, S. J. Martini coarse-grained force field: extension to DNA. Journal of chemical theory and computation 2015, 11, 3932–3945.

(65) Chakraborty, D.; Hori, N.; Thirumalai, D. Sequence-dependent three interaction site model for single-and double-stranded DNA. Journal of chemical theory and computation 2018, 14, 3763–3779.

(66) Snodin, B. E.; Schreck, J. S.; Romano, F.; Louis, A. A.; Doye, J. P. Coarse-grained modelling of the structural properties of DNA origami. Nucleic acids research 2019, 47, 1585–1597.

(67) Brandani, G. B.; Niina, T.; Tan, C.; Takada, S. DNA sliding in nucleosomes via twist defect propagation revealed by molecular simulations. Nucleic acids research 2018, 46, 2788–2801.

(68) Ouldridge, T. E.; Louis, A. A.; Doye, J. P. Structural, mechanical, and thermodynamic properties of a coarse-grained DNA model. The Journal of chemical physics 2011, 134, 02B627.

(69) Case, D. A.; Cheatham III, T. E.; Darden, T.; Gohlke, H.; Luo, R.; Merz Jr, K. M.; Onufriev, A.; Simmerling, C.; Wang, B.; Woods, R. J. The Amber biomolecular simulation programs. Journal of computational chemistry 2005, 26, 1668–1688.

(70) Hinckley, D. M.; Freeman, G. S.; Whitmer, J. K.; De Pablo, J. J. An experimentally-informed coarse-grained 3-site-per-nucleotide model of DNA: Structure, thermodynamics, and dynamics of hybridization. The Journal of chemical physics 2013, 139, 10B604_1.

(71) Olson, W. K.; Manning, G. S. A configurational interpretation of the axial phosphate spacing in polynucleotide helices and random coils. Biopolymers: Original Research on Biomolecules 1976, 15, 2391–2405.

(72) Assenza, S.; Mezzenga, R. Soft condensed matter physics of foods and macronutrients. Nature Reviews Physics 2019, 1, 551–566.

(73) Case, D. A.; Babin, V.; Berryman, J. T.; Betz, R. M.; Cai, Q.; Cerutti, D. S.; Cheatham III, T. E.; Darden, T. A.; Duke, R. E.; Gohlke, H. e. a. AMBER 14; University of California Press, 2014.

(74) Wang, J.; Cieplak, P.; Kollman, P. A. How well does a restrained electrostatic potential (RESP) model perform in calculating conformational energies of organic and biological molecules? Journal of computational chemistry 2000, 21, 1049–1074.

(75) Pérez, A.; Marchán, I.; Svozil, D.; Sponer, J.; Cheatham III, T. E.; Laughton, C. A.; Orozco, M. Refinement of the AMBER force field for nucleic acids: improving the description of *α/γ* conformers. Biophysical journal 2007, 92, 3817–3829.

(76) Galindo-Murillo, R.; Robertson, J. C.; Zgarbová, M.; Sponer, J.; Otyepka, M.; Jurecka, P.; Cheatham III, T. E. Assessing the current state of amber force field modifications for DNA. Journal of chemical theory and computation 2016, 12, 4114–4127.

(77) Ivani, I.; Dans, P. D.; Noy, A.; Pérez, A.; Faustino, I.; Hospital, A.; Walther, J.; Andrio, P.; Goñi, R.; Balaceanu, A., et al. Parmbsc1: a refined force field for DNA simulations. Nature methods 2016, 13, 55–58.

(78) Zgarbová, M.; Sponer, J.; Otyepka, M.; Cheatham III, T. E.; Galindo-Murillo, R.; Jurecka, P. Refinement of the sugar–phosphate backbone torsion beta for AMBER force fields improves the description of Z-and B-DNA. Journal of chemical theory and computation 2015, 11, 5723–5736.

(79) Plimpton, S. Fast parallel algorithms for short-range molecular dynamics. Journal of computational physics 1995, 117, 1–19.

(80) Snodin, B. E.; Randisi, F.; Mosayebi, M.; Sulc, P.; Schreck, J. S.; Romano, F.; Ouldridge, T. E.; Tsukanov, R.; Nir, E.; Louis, A. A.; Doye, J. P. Introducing improved structural properties and salt dependence into a coarse-grained model of DNA. The Journal of chemical physics 2015, 142, 06B613_1.

(81) Henrich, O.; Gutierrez-Fosado, Y. A.; Curk, T.; Ouldridge, T. E. Coarse-grained simulation of DNA using LAMMPS. The European Physical Journal E 2018, 41, 57.

(82) Freeman, G. S.; Hinckley, D. M.; Lequieu, J. P.; Whitmer, J. K.; De Pablo, J. J. Coarse-grained modeling of DNA curvature. The Journal of chemical physics 2014, 141, 10B615_1.

(83) Sassi, A. S.; Assenza, S.; De Los Rios, P. Shape of a stretched polymer. Physical review letters 2017, 119, 037801.

(84) Lu, X.-J.; Olson, W. K. 3DNA: a software package for the analysis, rebuilding and visualization of three-dimensional nucleic acid structures. Nucleic acids research 2003, 31, 5108–5121.

(85) Lavery, R.; Moakher, M.; Maddocks, J. H.; Petkeviciute, D.; Zakrzewska, K. Conformational analysis of nucleic acids revisited: Curves+. Nucleic acids research 2009, 37, 5917–5929.

(86) Yuksel, C.; Schaefer, S.; Keyser, J. Parameterization and applications of Catmull–Rom curves. Computer-Aided Design 2011, 43, 747–755.

(87) Trifonov, E. N.; Tan, R. K.-Z.; Harvey, S. Static persistence length of DNA. Structure and expression: proceedings of the Fifth Conversation in the Discipline Biomolecular Stereodynamics held at the State University of New York at Albany, June 2-6, 1987/edited by MH Sarma & RH Sarma 1988,

(88) Frenkel, D.; Smit, B. Understanding molecular simulation: from algorithms to applications; Elsevier, 2001; Vol. 1.

(89) Kabsch, W. A solution for the best rotation to relate two sets of vectors. Acta Crystallographica Section A: Crystal Physics, Diffraction, Theoretical and General Crystallography 1976, 32, 922–923.

(90) Rubinstein, M.; Colby, R. H., et al. Polymer physics; Oxford university press New York, 2003; Vol. 23.

(91) Vologodskaia, M.; Vologodskii, A. Contribution of the intrinsic curvature to measured DNA persistence length. Journal of molecular biology 2002, 317, 205–213.

(92) Moroz, J. D.; Nelson, P. Entropic elasticity of twist-storing polymers. Macromolecules 1998, 31, 6333–6347.

(93) Sheinin, M. Y.; Wang, M. D. Twist–stretch coupling and phase transition during DNA supercoiling. Physical Chemistry Chemical Physics 2009, 11, 4800–4803.

(94) Noy, A.; Golestanian, R. Length scale dependence of DNA mechanical properties. Physical review letters 2012, 109, 228101.

(95) Skoruppa, E.; Nomidis, S. K.; Marko, J. F.; Carlon, E. Bend-induced twist waves and the structure of nucleosomal DNA. Physical review letters 2018, 121, 088101.

(96) Nomidis, S. K.; Skoruppa, E.; Carlon, E.; Marko, J. F. Twist-bend coupling and the statistical mechanics of the twistable wormlike-chain model of DNA: Perturbation theory and beyond. Physical Review E 2019, 99, 032414.

