## Supplemental Information for "Accurate sequence-dependent coarse-grained model for conformational and elastic properties of double-stranded DNA"

### Supplementary Information

### S1 Coarse-graining procedure

In MADna, six different moieties are considered as effective particles, corresponding to the sugar, the phosphate group and the four bases. The effective particles are located in the geometrical centers of the heavy atoms belonging to each moiety. Specifically, for a sugar particle the center is computed considering the atoms C1', C2', C3', C4', C5' and O4', while for the phosphate group the atoms are O3', P, OP1, OP2 and O5. The rest of heavy atoms within the nucleotide are assigned to the base. The size $\sigma$ of each bead was fixed following Ref.[1]. The values are reported in Table S1.

| Name | Abbreviation | $\sigma$ (Å) | mass (g/mol) |
|---|---|---|---|
| Sugar | S | 6.4 | 76.05 |
| Phosphate group | P | 4.5 | 94.97 |
| Adenine | A | 5.4 | 130.09 |
| Cytosine | C | 6.4 | 106.06 |
| Guanine | G | 4.9 | 146.09 |
| Thymine | T | 7.1 | 120.07 |

Table S1: Excluded volume size ($\sigma$) and mass of the various beads.

### S2 Determination of the force-field parameters

The force-field parameters were fixed by matching coarse-grained predictions to the atomistic results obtained for the Learning Sequences. A Boltzmann Inversion was performed by considering the average and fluctuations size of the various bonded interactions for the all-atom simulations performed at 1 pN. More in detail, the values of $r_0$, $\theta_0$ and $\phi_0$ were fixed as the averages obtained from the coarse-grained trajectory of atomistic simulations (in the case of $\phi_0$, the average was shifted by $\pi$ since the minimum of the dihedral potential is found at $\phi_0 + \pi$). The elastic constants were tuned in order to match the size of fluctuations computed from the trajectories to the ensemble averages calculated from Equations (1), (2) and (3) in the main text, which we report here for convenience:

$$U_{\text{bond}}(r) = k_{\text{bond}}(r{-}r_0)^2 \,, \tag{1}$$

$$U_{\text{angle}}(\theta) = k_{\text{angle}}(\theta{-}\theta_0)^2 \,, \tag{2}$$

$$U_{\text{dihedral}}(\phi) = k_{\text{dihedral}}\left[1 + \cos(\phi - \phi_0)\right] \,. \tag{3}$$

In the case of bonds, the fluctuation size $\sigma_{\text{b}}$ was calculated as

$$\sigma_{\text{b}} = \sqrt{\frac{\int_0^\infty (r - r_0)^2 \cdot r^2 e^{-\frac{U_{\text{bond}}(r)}{k_B T}} dr}{\int_0^\infty r^2 e^{-\frac{U_{\text{bond}}(r)}{k_B T}} dr}} \,, \tag{S1}$$

with the weight factor $r^2$ being introduced to account for the three-dimensional nature of the system. Analogously, one has

$$\sigma_{\text{a}} = \sqrt{\frac{\int_0^\pi (\theta - \theta_0)^2 \cdot \sin\theta \, e^{-\frac{U_{angle}(\theta)}{k_B T}} d\theta}{\int_0^\pi \sin\theta \, e^{-\frac{U_{angle}(\theta)}{k_B T}} d\theta}} \tag{S2}$$

for the angles, while

$$\sigma_{\text{d}} = \sqrt{\frac{\int_0^{2\pi} (\phi - \phi_0 - \pi)^2 \cdot e^{-\frac{U_{dihedral(\phi)}}{k_B T}} d\phi}{\int_0^{2\pi} e^{-\frac{U_{dihedral(\phi)}}{k_B T}} d\phi}} \tag{S3}$$

holds in the case of dihedrals. As mentioned above, for each bonded interaction the mean and fluctuation size were computed from the coarse-grained atomistic trajectory and the equations relevant to the case at hand were employed to determine the force-field parameters. For instance, for a bond, Equation (S1) was inverted to determine the value of $k_{\text{bond}}$ from the knowledge of $\sigma_{\text{b}}$. This approach neglects the possible effects of the pulling force present in the atomistic simulations. In order to minimize this perturbation, the inversion procedure was

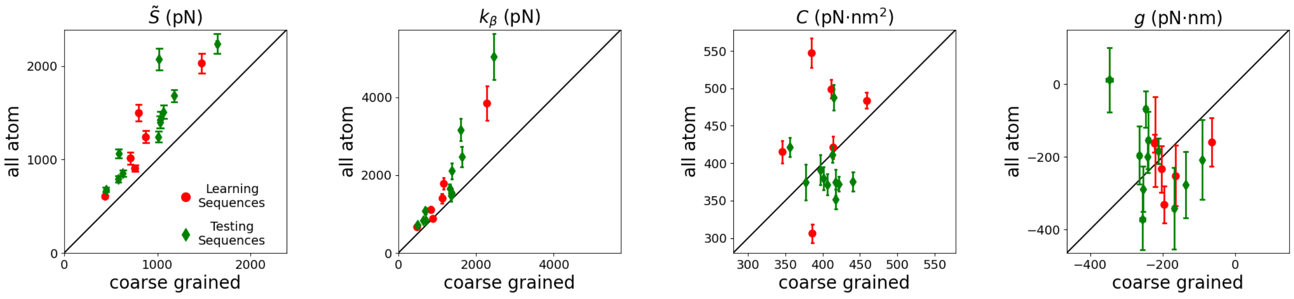


Figure S1: Scatter plot comparing atomistic and coarse-grained Boltzmann-Inversion results for the effective stretching modulus $\tilde{S}$, the crookedness rigidity $k_\beta$, the torsion modulus $C$ and the twist-stretch coupling constant $g$. Learning and Testing Sequences are denoted by red circles and green diamonds, respectively. The black line indicates the bisector of the first and third quadrant.

performed considering the atomistic trajectories obtained for the smallest pulling force, $f = 1$ pN.

With the force field obtained from Boltzmann Inversion, we performed coarse-grained pulling simulations and compared the obtained elastic constants with the atomistic results. It is evident from Fig.S1 that, while the coarse-grained model gives reasonable values, there is room for improvement. In order to obtain a better agreement, we thus proceeded to a refinement of the bonded parameters. The main goal of this second step was to reproduce the elastic constants of the Learning Sequences. In practice, this was done by iteratively adapting the parameters and running coarse-grained simulations of the Learning Sequences, from which the elastic constants were computed. Due to the large number of bonded parameters, an exhaustive search of the optimal set is beyond current computational possibilities. As discussed in the main text, we adjusted a minimal subset of key parameters which were observed to affect the elastic properties, namely the rigidities of bond 5'-BB-3', angle 5'-PSB-3' and dihedral 5'-SPSP-3'. In order to give an idea of the extent of the tuning of parameters, we mention that the most extreme changes were given by a factor equal to 0.25 (for angle 5'-PSB-3' and step AG) and a 3-fold increase (for dihedral 5'-SPSP-3' and step CG). The optimized set of parameters is reported in Tables S2, S3 and S4 for bonds, angles and dihedrals, respectively.

Bonds

| Step | 5'-SP-3' $r_0$ (Å) | 5'-SP-3' $k_{\text{bond}}$ $\left(\frac{\text{Kcal}}{\text{mol Å}^2}\right)$ | 5'-PS-3' $r_0$ (Å) | 5'-PS-3' $k_{\text{bond}}$ $\left(\frac{\text{Kcal}}{\text{mol Å}^2}\right)$ | 5'-BB-3' $r_0$ (Å) | 5'-BB-3' $k_{\text{bond}}$ $\left(\frac{\text{Kcal}}{\text{mol Å}^2}\right)$ |
|---|---|---|---|---|---|---|
| AA | 3.74 | 54.42 | 4.08 | 14.59 | 3.87 | 5.86 |
| AC | 3.74 | 58.05 | 4.07 | 15.03 | 3.73 | 8.02 |
| AG | 3.75 | 65.28 | 4.09 | 11.51 | 4.12 | 3.48 |
| AT | 3.75 | 67.15 | 4.12 | 15.87 | 3.70 | 18.57 |
| CA | 3.76 | 58.90 | 4.13 | 14.82 | 4.24 | 5.84 |
| CC | 3.77 | 79.36 | 4.17 | 19.01 | 4.19 | 2.28 |
| CG | 3.76 | 60.94 | 4.09 | 10.43 | 4.26 | 4.46 |
| CT | 3.75 | 75.03 | 4.12 | 18.69 | 3.95 | 4.00 |
| GA | 3.74 | 49.47 | 4.11 | 15.13 | 3.82 | 6.07 |
| GC | 3.71 | 41.33 | 4.04 | 19.21 | 3.71 | 9.52 |
| GG | 3.76 | 75.12 | 4.17 | 16.75 | 4.15 | 3.08 |
| GT | 3.74 | 54.90 | 4.11 | 17.11 | 3.69 | 13.19 |
| TA | 3.76 | 69.77 | 4.14 | 19.27 | 4.36 | 8.33 |
| TC | 3.76 | 73.87 | 4.17 | 19.81 | 4.19 | 2.96 |
| TG | 3.76 | 57.77 | 4.13 | 16.39 | 4.48 | 3.96 |
| TT | 3.76 | 79.95 | 4.14 | 23.67 | 4.02 | 4.66 |

| Base | SB $r_0$ (Å) | SB $k_{\text{bond}}$ $\left(\frac{\text{Kcal}}{\text{mol Å}^2}\right)$ |
|---|---|---|
| A | 4.89 | 39.23 |
| C | 4.39 | 44.51 |
| G | 5.01 | 49.13 |
| T | 4.46 | 45.25 |

| WC Pair | BB-WC $r_0$ (Å) | BB-WC $k_{\text{bond}}$ $\left(\frac{\text{Kcal}}{\text{mol Å}^2}\right)$ |
|---|---|---|
| AT | 6.09 | 21.48 |
| CG | 5.70 | 35.95 |

Table S2: Optimal parameter values for bonds (compare Equation (2)). Units are directly following the convention used in LAMMPS [4], when the option "units real" is selected.

Angles

| Step | 5'-SPS-3' $\theta_0$ (deg) | 5'-SPS-3' $k_{\text{angle}}$ $\left(\frac{\text{Kcal}}{\text{mol rad}^2}\right)$ | 3'-PSB-5' $\theta_0$ (deg) | 3'-PSB-5' $k_{\text{angle}}$ $\left(\frac{\text{Kcal}}{\text{mol rad}^2}\right)$ | 5'-PSB-3' $\theta_0$ (deg) | 5'-PSB-3' $k_{\text{angle}}$ $\left(\frac{\text{Kcal}}{\text{mol rad}^2}\right)$ |
|---|---|---|---|---|---|---|
| AA | 94.12 | 26.15 | 115.33 | 32.47 | 113.21 | 6.21 |
| AC | 91.81 | 27.73 | 116.09 | 40.51 | 109.29 | 20.74 |
| AG | 94.11 | 23.84 | 115.28 | 35.91 | 119.24 | 2.44 |
| AT | 92.83 | 33.04 | 114.73 | 43.58 | 105.79 | 20.24 |
| CA | 96.24 | 22.69 | 119.22 | 38.06 | 110.49 | 13.46 |
| CC | 94.52 | 32.47 | 117.56 | 51.38 | 111.33 | 19.19 |
| CG | 95.46 | 21.80 | 120.50 | 37.77 | 113.34 | 8.21 |
| CT | 92.87 | 38.18 | 117.21 | 51.27 | 106.69 | 4.26 |
| GA | 95.14 | 22.91 | 111.77 | 29.27 | 108.34 | 1.56 |
| GC | 92.66 | 25.46 | 111.94 | 33.19 | 106.68 | 10.49 |
| GG | 94.25 | 34.28 | 109.64 | 52.70 | 121.94 | 11.45 |
| GT | 94.46 | 28.77 | 111.28 | 37.18 | 104.88 | 26.48 |
| TA | 94.39 | 26.40 | 119.62 | 40.82 | 114.26 | 8.79 |
| TC | 93.81 | 34.18 | 121.01 | 48.59 | 107.49 | 3.25 |
| TG | 93.12 | 29.86 | 120.09 | 42.72 | 118.63 | 15.01 |
| TT | 92.66 | 46.62 | 119.89 | 57.92 | 104.95 | 13.15 |

| WC Pair | SBB $\theta_0$ (deg) | SBB $k_{\text{angle}}$ $\left(\frac{\text{Kcal}}{\text{mol rad}^2}\right)$ |
|---|---|---|
| AT | 154.31 | 51.58 |
| CG | 138.35 | 53.86 |
| GC | 158.75 | 52.15 |
| TA | 132.80 | 49.68 |

Table S3: Optimal parameter values for angles (compare Equation (3)). Note that, following the convention in LAMMPS [4], $\theta_0$ is expressed in degrees while $k_{\text{angle}}$ has units of Kcal/mol rad$^2$. Units are directly following the convention used in LAMMPS, when the option "units real" is selected. In the case of SBB, the Watson-Crick pairs are ordered by listing first the base bonded to the sugar moiety.

Dihedrals

| Step | 5'-SPSP-3' $\phi_0$ (deg) | 5'-SPSP-3' $k_{\text{dihedral}}$ $\left(\frac{\text{Kcal}}{\text{mol}}\right)$ | 5'-PSPS-3' $\phi_0$ (deg) | 5'-PSPS-3' $k_{\text{dihedral}}$ $\left(\frac{\text{Kcal}}{\text{mol}}\right)$ | 5'-SPSB-3' $\phi_0$ (deg) | 5'-SPSB-3' $k_{\text{dihedral}}$ $\left(\frac{\text{Kcal}}{\text{mol}}\right)$ |
|---|---|---|---|---|---|---|
| AA | 0.43 | 59.55 | 22.36 | 4.72 | -142.34 | 13.82 |
| AC | -1.50 | 23.66 | 27.39 | 4.82 | -141.29 | 19.09 |
| AG | -2.16 | 47.34 | 26.73 | 5.56 | -145.79 | 10.12 |
| AT | -0.87 | 19.10 | 24.42 | 7.50 | -137.42 | 27.14 |
| CA | -2.64 | 25.63 | 20.00 | 5.32 | -144.63 | 10.96 |
| CC | -3.69 | 44.82 | 23.61 | 11.51 | -140.36 | 21.61 |
| CG | -0.79 | 54.62 | 26.55 | 4.11 | -149.56 | 7.56 |
| CT | -0.50 | 57.43 | 24.68 | 10.46 | -136.72 | 27.53 |
| GA | -0.47 | 50.07 | 19.19 | 4.60 | -139.81 | 16.59 |
| GC | 0.98 | 52.40 | 17.71 | 2.75 | -137.94 | 21.41 |
| GG | -3.17 | 42.78 | 23.55 | 10.89 | -141.02 | 13.63 |
| GT | 0.70 | 25.74 | 19.03 | 5.61 | -135.20 | 28.05 |
| TA | -2.86 | 19.63 | 22.11 | 8.35 | -143.86 | 14.21 |
| TC | 0.07 | 60.41 | 19.32 | 10.76 | -138.01 | 23.55 |
| TG | -2.67 | 32.65 | 26.50 | 7.75 | -146.92 | 9.02 |
| TT | 2.02 | 77.55 | 20.52 | 15.66 | -135.61 | 34.02 |

| Step | 3'-SPSB-5' $\phi_0$ (deg) | 3'-SPSB-5' $k_{\text{dihedral}}$ $\left(\frac{\text{Kcal}}{\text{mol}}\right)$ | 5'-PSBB-3' $\phi_0$ (deg) | 5'-PSBB-3' $k_{\text{dihedral}}$ $\left(\frac{\text{Kcal}}{\text{mol}}\right)$ | 3'-PSBB-5' $\phi_0$ (deg) | 3'-PSBB-5' $k_{\text{dihedral}}$ $\left(\frac{\text{Kcal}}{\text{mol}}\right)$ |
|---|---|---|---|---|---|---|
| AA | 163.97 | 10.39 | 20.94 | 10.50 | -124.78 | 10.12 |
| AC | 166.22 | 10.91 | 19.79 | 13.64 | -120.21 | 7.45 |
| AG | 165.09 | 8.94 | 22.43 | 6.70 | -120.17 | 8.56 |
| AT | 164.45 | 14.86 | 21.44 | 23.90 | -116.14 | 8.18 |
| CA | 155.62 | 6.91 | 23.13 | 7.62 | -117.01 | 8.98 |
| CC | 157.37 | 13.92 | 8.61 | 17.91 | -126.38 | 13.92 |
| CG | 159.63 | 5.56 | 38.23 | 6.39 | -113.08 | 8.85 |
| CT | 158.95 | 13.43 | 21.42 | 18.04 | -119.92 | 10.04 |
| GA | 165.40 | 11.53 | 21.21 | 10.67 | -124.79 | 7.92 |
| GC | 165.80 | 11.04 | 23.86 | 15.31 | -114.10 | 6.71 |
| GG | 166.73 | 15.04 | 10.35 | 7.24 | -130.58 | 7.91 |
| GT | 165.12 | 14.54 | 22.28 | 24.12 | -113.48 | 7.56 |
| TA | 152.13 | 8.69 | 27.76 | 8.40 | -112.76 | 12.11 |
| TC | 149.37 | 12.01 | 19.82 | 12.94 | -112.15 | 12.99 |
| TG | 154.62 | 6.71 | 35.89 | 6.90 | -110.67 | 10.70 |
| TT | 151.88 | 15.72 | 21.43 | 21.30 | -115.81 | 14.63 |

Table S4: Optimal parameter values for dihedrals (compare Equation (4)). Units are directly following the convention used in LAMMPS [4], when the option “units real” is selected.

### S3 List of sequences

#### S3.1 Model definition and benchmark simulations

##### Learning Sequences

In order to determine the parameters of the bonded interactions, a matching procedure was employed to reproduce the results of a set of atomistic simulations exhaustively accounting for all the possible base steps [2]. This set is here referred to as "Learning Sequences" and is composed of the six molecules polyAA, polyAC, polyAG, polyAT, polyCG and polyGG, each consisting of a pentamer of dinucleotides flanked by two CGCG handles, introduced to avoid end effects. Hence, the sequences are (handles are evidenced by the presence of a dash in the sequence):

polyAA ↔ 5'-CGCG-AAAAAAAAAA-CGCG-3',

polyAC ↔ 5'-CGCG-ACACACACAC-CGCG-3',

polyAG ↔ 5'-CGCG-AGAGAGAGAG-CGCG-3',

polyAT ↔ 5'-CGCG-ATATATATAT-CGCG-3',

polyCG ↔ 5'-CGCG-CGCGCGCGCG-CGCG-3',

polyGG ↔ 5'-CGCG-GGGGGGGGGG-CGCG-3'.

##### Testing Sequences

The predictive power of the model was tested by comparing the coarse-grained and atomistic results for a second set of sequences [2, 3], which are referred to as "Testing Sequences". Some of these sequences have direct biological relevance, such as the DNA Unwinding Element (DUE), a TATA-box element (TATA) and a Transcription Factor Binding Site (TFBS). As before, the handles are evidenced by the presence of a dash in each sequence.

DDD ↔ 5'-CGCG-CGCGAATTCGCG-CGCG-3',

TATA ↔ 5'-CGCG-TATAAAAG-CGCG-3',

TFBS ↔ 5'-CGCG-GGATGGGAG-CGCG-3',

G4CG4 ↔ 5'-CGCG-GGGGCGGGG-CGCG-3',

G4AAG4 ↔ 5'-CGCG-GGGGAAGGGG-CGCG-3',

DUE ↔ 5'-CGCG-GATCTATTTATTT-CGCG-3',

A4TA4 ↔ 5'-CGCG-AAAATAAAA-CGCG-3',

A4GGA4 ↔ 5'-CGCG-AAAAGGAAAA-CGCG-3',

A8T ↔ 5'-CGCG-AAAAAAAAT-CGCG-3',

A8GG ↔ 5'-CGCG-AAAAAAAAGG-CGCG-3',

All-steps ↔ 5'-GCG-CAATGGAGTA-CGC-3'.

#### S3.2 Persistence-length simulations

**Sequence-averaged**

P1 ↔ 5'-ACTCATCGAC TGTATTGTGC TCGGTAGATT GAAGGAGTAG CTTAGTCCTC ATAAAGCGGA CGAACATCGT TGCTAGAAAA TCCCTGGAAT GCACGTAGAT-3'

P2 ↔ 5'-GCAACAAGGT ATCAAGTTAT GCAGGGTCTT TCTGGGTTGT CGTACTAGCT GGATTAAATG CGAAGAGTCG CTAAAACCAC ACTACCAAAC ACTAGTAATA-3'

P3 ↔ 5'-CTAGACATTA TGGTTGGTGA GACCCGAGCC TTCTCTAAGA GACAACGCTA ATGTCTCGGA CGGTAGTCCG TCCCCTGAAT ATGCAATACA CCGTCTCTGA-3'

P4 ↔ 5'-GTCGTTTTCA CTCCCAGACC CATATGTTAT GTATTGTCCC AGTAATGTGT TGGTGCTAGA GCTAGCCTAT GTTCCAACCA CCAGCCTTCT GACTTTACAA-3'

P5 ↔ 5'-ACTTCACTTG TCGAAGTATT ACCTCATTAG AATAAATATT TTTATTCCAA GTCTCATCAT GGGGAAGGTG CTCTCATCTC TTTGCTGCGT AGTTTTATGG-3'

P6 ↔ 5'-GTTTGGTGCT TAGACCCGAT GCAACGGGCA TCATTACCCG GGTCCCCAAG GCCTAAGTAT GAATGTGTTA GACCCCCATC TTAGAACTGC TGGCGGCAAT-3'

P7 ↔ 5'-ATCGCGGGTT TAGGCTCGTC TCATTAGTTA AATTATGTGC GTCATAGAGA AGGTTTGAGC CTCGCATGAA GCTGTCAACG CCGATGTAGT TGTGTACGTG-3'

P8 ↔ 5'-GACTAACTAA GCACTGGGGA GTAACTGCTC CAGACTGTTA GACCTCTCAC TCCTAAAGGT ATTGAGAAAG CGTACTTCTA CATTCATTTG CACTGGTATC-3'

P9 ↔ 5'-TACCCGATAC GCTATCAGAA AGCATGGGCT AGTAAGCACA ATGGTACTGA GTACTCTACG TACGCAACCG TCCTTATTTA AATGCGCGGC CAAAACCAAA-3'

P10 ↔ 5'-CCAGGCGCCC GACTGCGATG GGCGATGCAA CTCCGAGTAG TTTAATTCGA GGGGAGCTGG AGGCATTGTA TCAGATTTTG AGCGGTGGCT TTTACCAGAG-3'

P11 ↔ 5'-GAGCATGCCG ATGTCCTGGC GACACGAAAC CCCCGAGCGC AACGCTGCTC ACAATGTTTT AGTATGATCC GACGTATGTA AAACGTCGTT GGGACCATTG-3'

P12 ↔ 5'-CACTTCAGGC GTCCTCTCCG CGGAACACTA GTACTCCCGT TCGGATGCTG GGGCTCCTTC TGTGAGGATT CAAAATGGGA AAGATCCCAA CCGGTGTGCC-3'

P13 ↔ 5'-TATAGGTCTG CGGCTCCCTT CTGTCAACTT GTCCTGCCAT CTCCCAAGTC GCTACGTGAG TTAGTCCCTT TAGCTGGAGC GAGGCATCCC TACAAGCCTG-3'

P14 ↔ 5'-ACGTATCGTG ACGAGATAGT CCTACTACTA GGAAGAAATC CCGGCTAGGT ATAAGAGGCC TGACCTAAAT CAAGAACAGA TGGGTTCAAT GAACTCATAG-3'

P15 ↔ 5'-CAAACCGCCT TGGGCTCGAG CCTTCGGGCG TGACGGGACG AGTCAATTTC AGTCCATACG CTAGTTATGG TATACAGGCG TCGGAGTTGC CCTTGATGGA-3'

P16 ↔ 5'-GGTTGCGATG AAAAGCTAGT GGTATTTGAC AAGCAGGCTT CAGTCACCCT GCGGCGTAGT TTTGTTAGAC ACTTCCCGTC TAGCTGTCGT ACATTTACGT-3'

P17 ↔ 5'-GTCTTTCGGG GGACGATAGG CGATCCCAAC TCTCTCTATC TAAGACGAAA TCAGTGCAAA AGTAATAGGG ATAACGTCCG CGTGTGTCAT TCGCCGTATC-3'

P18 ↔ 5'-GCCAGGCGGC GGTACATGGA GGATGCAAGA GTCTTATTGT GTTGTATAAG GCTTCACAGG TACCTATGCA TAGGCGGCTA AACTCTCACC CATGCCGACA-3'

P19 ↔ 5'-TTGAGAGATC GCCACACACT CAAGAGAGTG GACTCATGAA ACTTCCGACG TACGGACGCC GGATGCGGGG TATTATGGGC TGTAGCCGTA TGCGTGCCGC-3'

P20 ↔ 5'-CCGATCACAT GTCCGGCGCA ATCGGTATTT TGTGGGTCTA CAGGTACTAT GTTATTCGTA CGTTCTGGCC CCGATCACGA ACCGATGAGC ACAGATGCTG-3'

**Sequence-dependent**

ACAT ↔ 5'-TACACATATA TACACATACA TACACACATA CATATATATA CACACACATA TACATACATA TACATACATA CATATATACA CACATACATA CACATATATA-3'

ACCAGG ↔ 5'-AGGACCACCA GGAGGACCAG GAGGAGGACC ACCACCAGGA CCACCAGGAC CAGGAGGAGG ACCACCAGGA CCACCAGGAG GAGGACCACC AGGACCAGGA-3'

ACGAGC ↔ 5'-CACGACGACG AGCACGAGCA GCACGACGAG CACGAGCAGC AGCACGAGCA CGACGAGCAG CACGACGAGC AGCACGACGA GCACGACGAC GAGCAGCAGC-3'

AGAT ↔ 5'-AGAGATATAT AGAGATAGAT AGAGAGATAG ATATATATAG AGAGAGATAT AGATAGATAT AGATAGATAG ATATATAGAG AGATAGATAG AGATATATAT-3'

AGC ↔ 5'-GCAGCTAGCA GCAGCAGCAG CAAGCAGCAG CAGCAGCGAG CAGCAGCAGC AGCTAGCAGC AGCAGCAGCA AGCAGCAGCA GCAGCGAGCA GCAGCAGCAG-3'

CAA ↔ 5'-CAACAACAAG AACAACAACA ACAACAACCA ACAACAACAA CAACAACCAA CAACAACAAG CAACAACAAC AACAACAACC AACAACAACA ACCAACAACA-3'

CAACTT ↔ 5'-CTTCAACAAC TTCAACTTCT TCAACAACAA CTTCTTCTTC AACAACTTC[illegible] ACAACTTCTT CAACTTCAAC AACTTCTTCA ACTTCAACTT CTTCAACAAC-3'

CAGT ↔ 5'-AGCAGTCAGT CAGTCAGTCA GTCAGTCAGT CTGACAGTCA GTCAGTCAGT CAGTCAGTCA GTCAGTCTGA CAGTCAGTCA GTCAGTCAGT CAGTCAGTCA-3'

CATCTA ↔ 5'-ATCTACTACA TCTACTACTA CATCATCATC TACATCTACT ACATCATCTA CTACATCATC TACTACATCA TCTACATCTA CATCTACTAC ATCATCTACT-3'

HPL1 ↔ 5'-AGCGATTCGG CATTCGATTC GCGCATTCGA TTCGCATTCG ATTCGGCATT CGATTCGGCA TTCGATTCGC ATTCGATTCA TTCATTCGAT TCGGCATTCG-3'

HPL2 ↔ 5'-ACGACGAACG ACGAACGACG ACGAACGAAC GACGACGAAC GACGAACGAC GACGAACGAC GACGAACGAC GAGACGAACG AACGACGACG AACGACGAAC-3'

LPL1 ↔ 5'-GCAGTAGTAG CCTAGTAGCC TAGTAGCCTA GTAGCCTAGT AGCCTAGTAG CCTAGTAGCC TAGTAGCCTA GTAGCCTAGT AGCCTAGTAG CCTAGTAGCC-3'

LPL2 ↔ 5'-AGGGCCATAG GCATGCATAG GCATAGGCCA TGGCATAGGC ATTAGGCATG CATAGGCATA GGCATGGCAT AGGCATTAGG CATGCATAGG CATAGGCATG-3'

SG1 ↔ 5'-AGCAGCTAGC TAGCGCGATG CCCAGCTGAG ATCGACGATC GATGGCGATT ATCAGCTAGC AGCTAGCGAT CGACGCGCGA TGCGCAGCTG AGCTAGCTGA-3'

#### S3.3 Stretch-torsion simulations

**Sequence-averaged**

ST1 ↔ 5'-GATTAATGAC GGACAACTCT GCTGTCCTGA CGCGGCGAAA-3'

ST2 ↔ 5'-CGAATGGACG TGGAGGTAGT CCGATTTCGA TCCAAGGTGA-3'

ST3 ↔ 5'-TTCTCCTGCA ACTGGATCCC GCTTGATGTT TTAGGTTAAC-3'

ST4 ↔ 5'-TTCTAACTGA TTTGGCTAGC GTATACTTCT GTATCTGTCC-3'

ST5 ↔ 5'-CCTCACATTT TAGCACCGCC TCCCGATAAC TTAAGGGTTC-3'

ST1-short ↔ 5'-GGACAACTCT GCTGTCCTGA-3'

ST2-short ↔ 5'-TGGAGGTAGT CCGATTTCGA-3'

ST3-short ↔ 5'-ACTGGATCCC GCTTGATGTT-3'

ST4-short ↔ 5'-TTTGGCTAGC GTATACTTCT-3'

ST5-short ↔ 5'-TAGCACCGCC TCCCGATAAC-3'

**Phased A-tracts**

A-tract-1 ↔ 5'-CCCCAAAAAT TTTCAAATTT TTTCTCGAAA AATTGCAAAA-3'

A-tract-2 ↔ 5'-CAAATAATCA TTCAAAAAAA AAAACTTTCT AAAAAATCTC-3'

A-tract-3 ↔ 5'-CAAAAATTTT CAAATTTTTT CTCGAAAAAT TGCAAAAAAT-3'

A-tract-4 ↔ 5'-GAAAAAAAAT GCTTAAAATT TCAGAAAATG TTAAAAATTC-3'

A-tract-5 ↔ 5'-AAAAATTCCG AAAATTGGTT AAAAAATTTT TTTAAAAAAC-3'

#### S3.4 Twist-bend simulations

TB1 ↔ 5'-ACAGAACTAG TCGCTACGTA CATGGTCCAT CGCTGGTATA TGCATAATAT AGTGTGAGAA ACCCAATGGG CTATGCGCTC TTTGGGCCAC AGCCCCTGCA AAAATACTTG TGTACTACCT TCTCCTCAGC TGTCAGAGAA TACATGATCG-3'

TB2 ↔ 5'-TTCTAGGAGA CATGTATATC GGATATGTCA TTGTATTAGC GAAGAGCAGG AGGAAACGCT GCTCCTGTAA CAGGGTTGGT TCGGTGCAGA GGGGTAATTT GGACTCCAGC CCTACTGTCA AGTAACTAGG TTGCACAGAC CATTGAGAAC-3'

TB3 ↔ 5'-GTAAAGCGAT AGCAAGATGA CCTATAGTGA TCTTAAACGT CCACAGAGAT CAGTTTTTCC TCTCATATTC AGGGTCCGTA GAGATACGGG AGGGGATTAC TCAACAGT[illegible] GTGGGGGGGA TATATAACTC TCTTCTTGTC CGGTCGATGA-3'

### S4 Determination of helical parameters

#### S4.1 Best fitting cylinder

The helical radius and axis were defined by considering the cylinder which best fitted the positions of the phosphates on the two strands (see Fig.S2a). The fit was performed by implementing the algorithm proposed in Ref.[6], where the optimal axis is found by iteratively proposing random orientations and selecting the one minimizing the square error. For the selected axis, the radius is obtained as a byproduct of the computation.

Here we report for convenience the main steps of the derivation in Ref.[6]. An infinite cylinder is identified by a unit-length axis $\hat{\boldsymbol{H}} \equiv (h_0, h_1, h_2)$, a radius $R$ and a center $\boldsymbol{C}$ chosen arbitrarily along the axis. By introducing the identity matrix $I$ and the transposed vector $\hat{\boldsymbol{H}}^T$, the rejection operator $P \equiv I - \hat{\boldsymbol{H}}\hat{\boldsymbol{H}}^T$ projects any vector onto the plane perpendicular to $\hat{\boldsymbol{H}}$. By construction, one has $P^T = P$ and $P^2 = P$. For any point $\boldsymbol{X}$, the quantity $(\boldsymbol{X} - \boldsymbol{C})^T P^T P (\boldsymbol{X} - \boldsymbol{C}) = (\boldsymbol{X} - \boldsymbol{C})^T P (\boldsymbol{X} - \boldsymbol{C})$ corresponds to the square of the distance separating $\boldsymbol{X}$ from the axis. Hence, if $\boldsymbol{X}$ lies on the surface of the cylinder, the following formula holds:

$$(\boldsymbol{X} - \boldsymbol{C})^T P (\boldsymbol{X} - \boldsymbol{C}) = R^2 . \tag{S4}$$

For fitting purposes, the error function to be minimized is thus

$$E\left(R^2, \boldsymbol{C}, \hat{\boldsymbol{H}}\right) \equiv \sum_{i=1}^{n} \left[(\boldsymbol{X}_i - \boldsymbol{C})^T P (\boldsymbol{X}_i - \boldsymbol{C}) - R^2\right]^2 , \tag{S5}$$

where $\boldsymbol{X}_i, i = 1, \ldots, n$ are the points to be fitted. The best-fitting cylinder is found by minimizing the error function with respect to the radius, center and axis of the cylinder. Denoting as $R_i$ the distance separating $\boldsymbol{X}_i$ from the axis, i.e. $R_i^2 = (\boldsymbol{C} - \boldsymbol{X}_i)^T P (\boldsymbol{C} - \boldsymbol{X}_i)$, the condition $\partial E / \partial R^2 = 0$ implies

$$R^2 = \frac{1}{n} \sum_{i=1}^{n} R_i^2 . \tag{S6}$$

The condition $\partial E / \partial \boldsymbol{C} = 0$ provides an equation which does not uniquely identify $\boldsymbol{C}$, since any point along the axis is an acceptable solution. Choosing for the center the point satisfying the supplementary condition $P\boldsymbol{C} = \boldsymbol{C}$, it can be shown [6] that

$$\boldsymbol{C} = \frac{\bar{A}}{\text{Trace}\left(\bar{A}A\right)} \left[\frac{1}{n} \sum_{i=1}^{n} \left(\boldsymbol{X}_i^T P \boldsymbol{X}_i\right) \boldsymbol{X}_i\right] , \tag{S7}$$

where

$$A \equiv P \left(\frac{1}{n} \sum_{i=1}^{n} \boldsymbol{X}_i \boldsymbol{X}_i^T\right) P \tag{S8}$$

and

$$\bar{A} \equiv SAS^T \,, \tag{S9}$$

with $S$ being the skew symmetric matrix

$$S = \begin{pmatrix} 0 & -h_2 & h_1 \\ h_2 & 0 & -h_0 \\ -h_1 & h_0 & 0 \end{pmatrix} \,. \tag{S10}$$

The equation stemming from the condition $\partial E/\partial \hat{\boldsymbol{H}} = 0$ cannot be solved analytically, thus preventing the possibility of building a closed-form solution for the direction of the optimal cylinder. Nevertheless, for any given axis $\hat{\boldsymbol{H}}$, the optimal radius and center are readily found by means of Equations (S6) and (S7). Substitution of these values into Equation (S5) leads to a conditional least-square error $E_{\hat{\boldsymbol{H}}}$, i.e. the minimum error associated to the chosen axis $\hat{\boldsymbol{H}}$. Hence, the fit can be performed by considering many randomly-oriented trial axes and selecting the one with the lowest value of $E_{\hat{\boldsymbol{H}}}$.

In the present case, $10^4$ iterations were considered to search for the optimal cylindrical direction. We have counterchecked numerically the efficacy of this approach by extracting coordinate points from artificially-created double helices with known axes and radii, chosen at random within the typical ranges found for DNA. More in detail, we built $10^4$ double helices with orientation $\hat{\boldsymbol{h}}$ chosen at random. The radius $r$ and periodicity $l$ were also chosen at random within the intervals $[0.5 \text{ nm}, 5 \text{ nm}]$ and $[1 \text{ nm}, 10 \text{ nm}]$, respectively, which largely encompass the ranges of values typical of double-stranded DNA. The two helices were shifted in order to form a minor and major groove with sizes following the ratio 4:7, as found in the crystal structure of the Drew-Dickerson dodecamer [7]. For each helix, twenty equally-spaced points were extracted within two periodicities, mimicking the positions of the phosphate groups along DNA fragments of length comparable to the Learning and Testing Sequences considered in this work. For each helix, the search for the optimal axis was run by considering $10^4$ random orientations. The radius $R$ and axis $\hat{\boldsymbol{H}}$ of the fitted cylinder were then compared to $r$ and $\hat{\boldsymbol{h}}$. As for the radii, we considered the relative error $e_R \equiv |R - r|/r$. On average, it was found that $\langle e_R \rangle \simeq 0.002$. Moreover, its maximum value was 0.05 and an error $e_R > 0.01$ was found in only about 1% of the simulated helices. As for the orientation, we computed the angle $\theta$ formed by $\hat{\boldsymbol{h}}$ and $\hat{\boldsymbol{H}}$, i.e. $\cos\theta \equiv \hat{\boldsymbol{h}} \cdot \hat{\boldsymbol{H}}$. On average, we found that $\langle \theta \rangle \simeq 0.7$ degrees, with a maximum value of 2.6 degrees and $\theta > 1.5$ degrees found in about 4% of cases. Overall, these results corroborate the robustness of the algorithm for the determination of the helical features of double-stranded DNA.

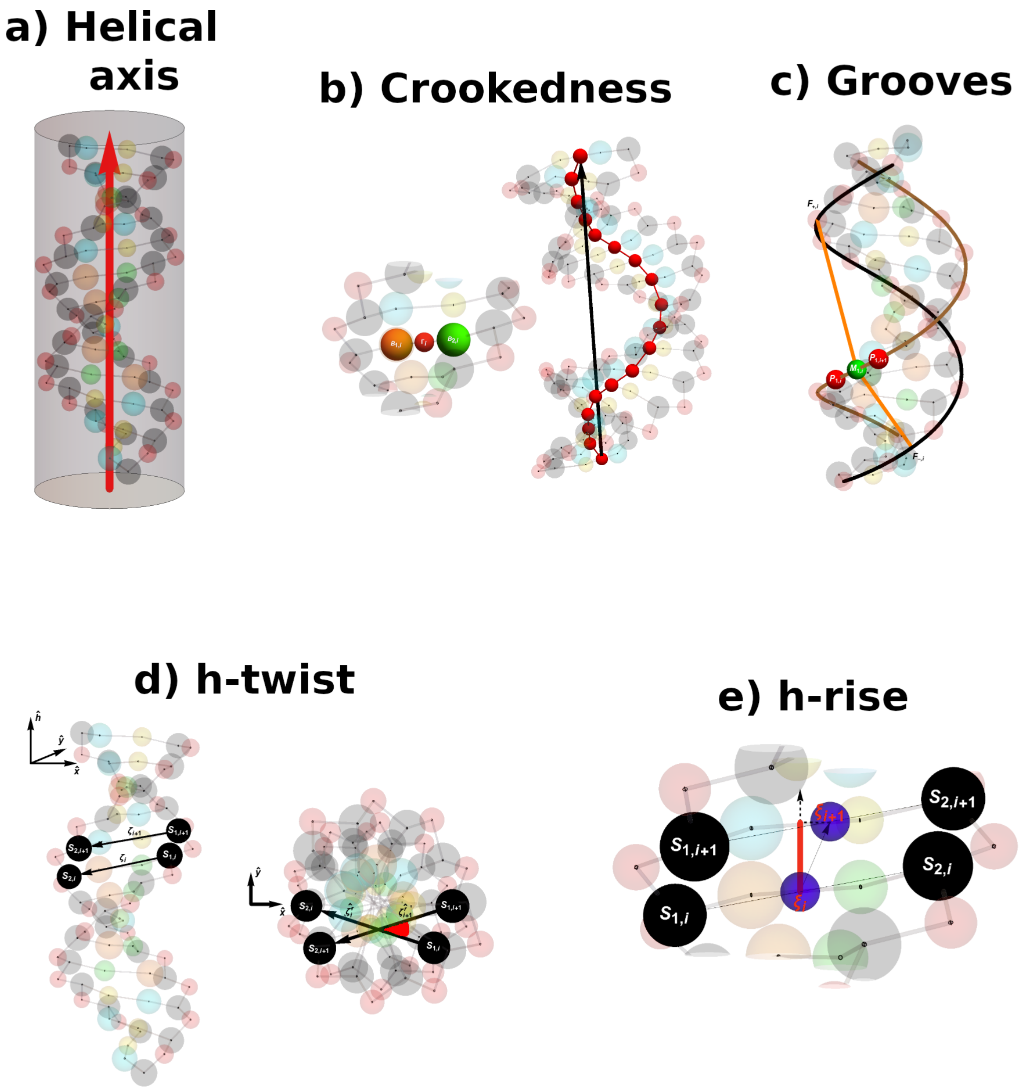


Figure S2: Sketches showing pictorially the various geometrical definitions reported in the Methods in the main text and in Section S4. a) The helical axis is obtained as the axis of the best-fitting cylinder. b) The crookedness is obtained by computing the arccosine of the ratio between the end-to-end distance (black arrow) and the contour-length of DNA (red line). These quantities are obtained along the line formed by the points $\Gamma_i$, which are determined as the centers between bases belonging to the same pair (inset). c) Grooves are defined by considering the lines interpolating the phospate beads (brown and black lines). For any couple of phosphates on the first strand ($\boldsymbol{P}_{1,i}$ and $\boldsymbol{P}_{1,i+1}$ in this case) we define the midpoint ($\boldsymbol{M}_i$). From the midpoint, we find the closest points on the second strand. The groove widths are obtained as the corresponding minimum distances (orange segments), suitably shifted to account for the excluded volume of the backbone. d) The h-twist is defined by considering the vectors $\boldsymbol{\zeta}_i \equiv \boldsymbol{S}_{2,i} - \boldsymbol{S}_{1,i}$ joining the two sugars within each base. The vectors $\boldsymbol{\zeta}_i$ are projected onto the plane perpendicular to the helical axis, thus obtaining $\boldsymbol{\zeta}_i^r$. The h-twist is then defined as the angle depicted in red, corresponding to cos h-twist $= \boldsymbol{\zeta}_i^r \cdot \boldsymbol{\zeta}_{i+1}^r$. e) The hrise is defined by considering the geometrical centers of the sugars $\xi_i \equiv (\boldsymbol{S}_{1,i} + \boldsymbol{S}_{2,i})/2$ and projecting the vector separating two consecutive centers onto the helical axis, thus obtaining h-rise $= (\xi_{i+1} - \xi_i) \cdot \hat{\boldsymbol{h}}$, corresponding to the red segment in the figure.

##### S4.2 Comparison between computed h-twist and 3DNA definition

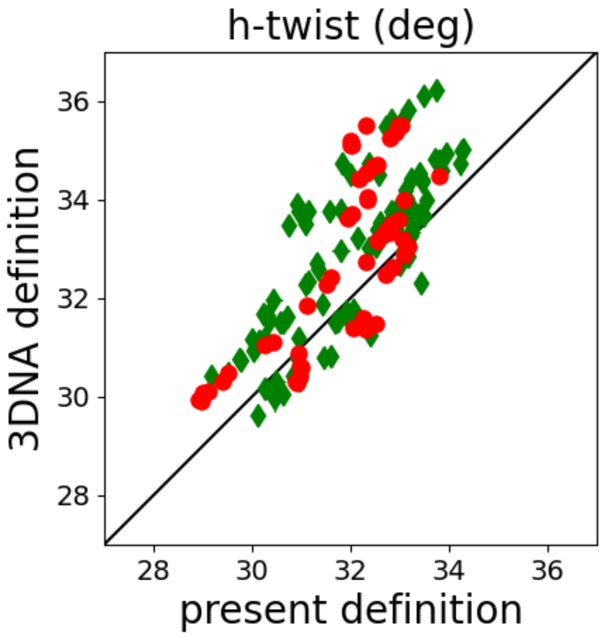


Figure S3: Scatter plot comparing the h-twist computed for the atomistic trajectories according to the present definition and the values obtained by means of 3DNA [8]. Red circles and green diamonds represent steps from Learning and Testing Sequences, respectively. The black line depicts the bisector of the first and third quadrant.

### S5 Benchmark Simulations

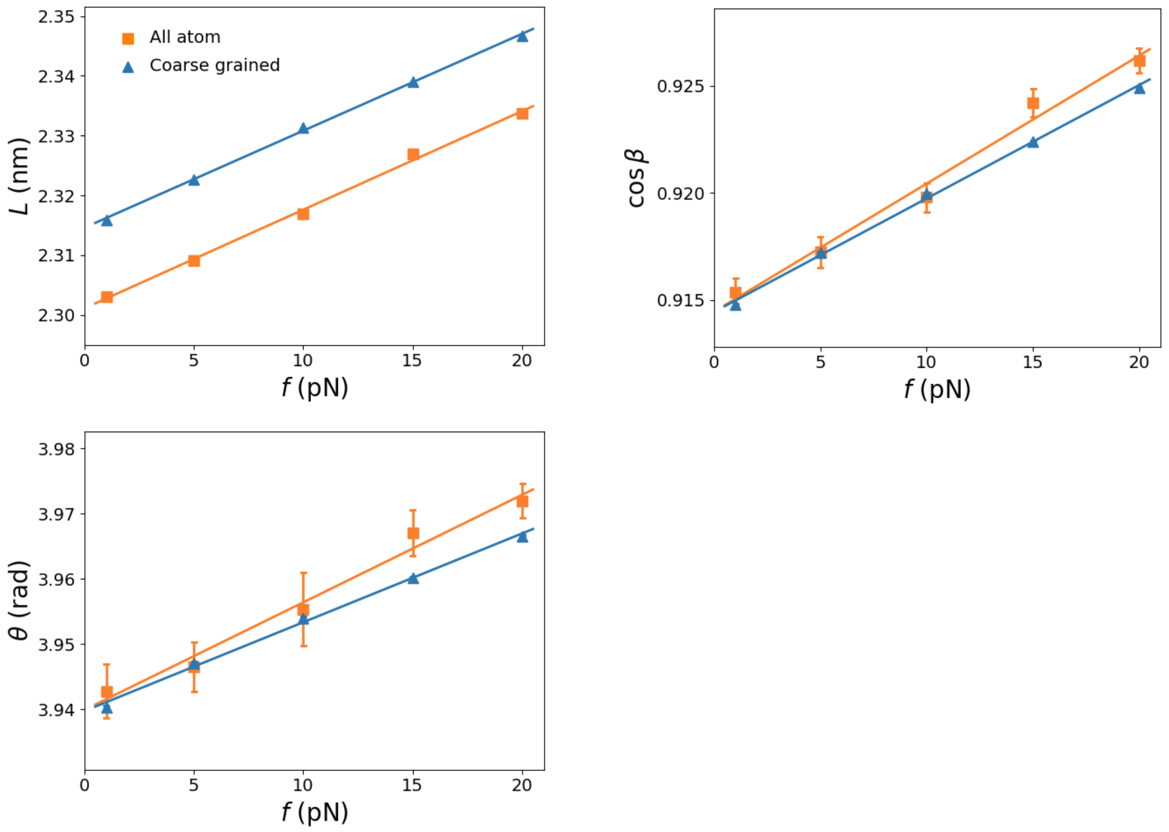


Figure S4: Linear dependence of extension $L$ (top-left panel), cosine of crookedness $\cos\beta$ (top-right panel) and cumulate h-twist $\theta$ (bottom-left panel) as a function of the force $f$ for a representative sequence (TATA). All-atom and coarse-grained results are denoted by orange squares and blue triangles, respectively. The lines show the linear fits of the data.

Learning Sequences

| Name | Simulation Type | $\tilde{S}$ (pN) | $k_\beta$ (pN) | $C$ (pN·nm$^2$) | $g$ (pN·nm) |
|---|---|---|---|---|---|
| AA | coarse grained | 2080 ± 15 | 3706 ± 37 | 419 ± 2 | -162 ± 8 |
| | all atom | 2031 ± 105 | 3848 ± 444 | 421 ± 15 | -159 ± 124 |
| AC | coarse grained | 972 ± 5 | 1058 ± 6 | 523 ± 3 | -257 ± 6 |
| | all atom | 1011 ± 65 | 1116 ± 73 | 548 ± 20 | -252 ± 83 |
| AG | coarse grained | 1163 ± 6 | 1460 ± 9 | 289 ± 2 | -230 ± 5 |
| | all atom | 1240 ± 61 | 1400 ± 120 | 306 ± 12 | -234 ± 64 |
| AT | coarse grained | 899 ± 5 | 897 ± 5 | 479 ± 2 | -331 ± 6 |
| | all atom | 905 ± 35 | 878 ± 46 | 499 ± 12 | -331 ± 50 |
| CG | coarse grained | 1442 ± 10 | 1859 ± 15 | 410 ± 2 | -159 ± 6 |
| | all atom | 1494 ± 87 | 1785 ± 151 | 415 ± 15 | -159 ± 67 |
| GG | coarse grained | 575 ± 2 | 671 ± 3 | 492 ± 3 | -175 ± 4 |
| | all atom | 605 ± 15 | 675 ± 21 | 484 ± 11 | -164 ± 24 |

Table S5: Comparison between atomistic and coarse-grained results for the Learning Sequences. The elastic constants were computed discarding the handles from the analysis.

Testing Sequences

| Name | Simulation Type | $\tilde{S}$ (pN) | $k_\beta$ (pN) | $C$ (pN·nm$^2$) | $g$ (pN·nm) |
|---|---|---|---|---|---|
| DDD | coarse grained | 1785 ± 14 | 2753 ± 29 | 372 ± 2 | -124 ± 6 |
| | all atom | 2070 ± 113 | 2475 ± 265 | 422 ± 13 | -208 ± 110 |
| TATA | coarse grained | 1427 ± 10 | 1727 ± 12 | 420 ± 2 | -353 ± 9 |
| | all atom | 1397 ± 65 | 1536 ± 113 | 371 ± 14 | -372 ± 83 |
| TFBS | coarse grained | 781 ± 5 | 891 ± 5 | 388 ± 2 | -243 ± 6 |
| | all atom | 851 ± 34 | 832 ± 45 | 380 ± 15 | -289 ± 62 |
| G4CG4 | coarse grained | 647 ± 4 | 741 ± 4 | 466 ± 3 | -211 ± 5 |
| | all atom | 678 ± 25 | 726 ± 37 | 372 ± 10 | -184 ± 35 |
| G4AAG4 | coarse grained | 773 ± 3 | 933 ± 5 | 424 ± 2 | -216 ± 4 |
| | all atom | 791 ± 33 | 837 ± 45 | 375 ± 13 | -199 ± 45 |
| DUE | coarse grained | 1412 ± 10 | 1740 ± 15 | 380 ± 2 | -217 ± 6 |
| | all atom | 1421 ± 91 | 1466 ± 148 | 391 ± 20 | -342 ± 113 |
| A4TA4 | coarse grained | 1431 ± 14 | 1790 ± 15 | 425 ± 2 | -311 ± 9 |
| | all atom | 1508 ± 71 | 2103 ± 197 | 374 ± 18 | -196 ± 80 |
| A4GGA4 | coarse grained | 1402 ± 8 | 1967 ± 13 | 388 ± 2 | -215 ± 6 |
| | all atom | 1242 ± 57 | 1648 ± 116 | 352 ± 13 | -155 ± 79 |
| A8T | coarse grained | 2303 ± 33 | 3741 ± 46 | 427 ± 2 | -298 ± 13 |
| | all atom | 2235 ± 102 | 5039 ± 582 | 488 ± 17 | 12 ± 89 |
| A8GG | coarse grained | 1629 ± 10 | 2532 ± 20 | 410 ± 2 | -197 ± 7 |
| | all atom | 1681 ± 65 | 3166 ± 285 | 411 ± 11 | -69 ± 50 |
| All-steps | coarse grained | 785 ± 3 | 880 ± 4 | 396 ± 2 | -193 ± 4 |
| | all atom | 1064 ± 48 | 1080 ± 86 | 374 ± 24 | -277 ± 91 |

Table S6: Comparison between atomistic and coarse-grained results for the Testing Sequences. The elastic constants were computed discarding the handles from the analysis.

#### S5.1 Analysis of A8T

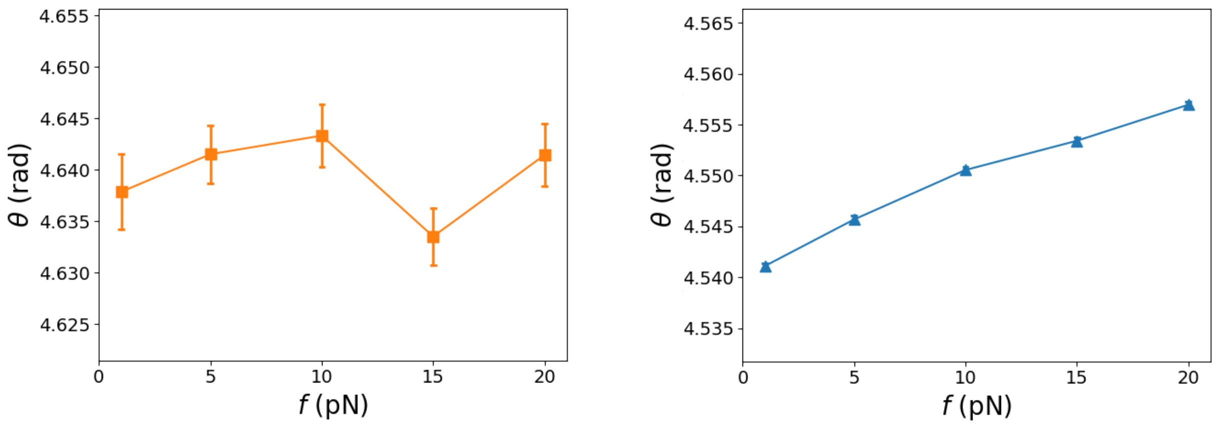


Figure S5: Dependence of cumulate h-twist $\theta$ on the force $f$ for sequence A8T. Left and right panel correspond to all-atom and coarse-grained results, respectively.

As mentioned in the main text, in the case of sequence A8T a discrepancy is observed between the coarse-grained and atomistic predictions. The coarse-grained model predicts $g = (-298 \pm 13)$ pN·nm, while $g = (12 \pm 89)$ pN·nm according to atomistic simulations (cfr Table S6). This discrepancy can be ascribed to the noisy response to the pulling force observed in the latter case for large values of $f$ (Fig.S5 left). Indeed, if one limits the analysis to low forces, $f \leq 10$ pN, the atomistic prediction becomes $g = (-242 \pm 237)$ pN·nm, which is in agreement with the coarse-grained prediction, although being characterized by a large error. The flucutating behavior of the cumulate h-twist observed for atomistic simulations is likely due to a lack of full convergence at large forces.

### S6 Robustness of persistence-length results with respect to definition of tangent vectors

The estimation of the persistence length of dsDNA may be sensitive to the choice made for the tangent vectors $\hat{\boldsymbol{t}}_i$ [5]. In order to assess the robustness of our results, we considered four different definitions for the tangent vectors, to which we refer as Sugars, Cylinder, Bases and Helical axis definitions.

The Sugars definition corresponds to the one reported in the main text. For the Cylinder definition, we define the displacement vector as $\boldsymbol{R}_{ij} \equiv \boldsymbol{O}_j - \boldsymbol{O}_i$, where $\boldsymbol{O}_i$ is the centre of the cylinder defining the helical axis in the fragment between $i-5$ and $i+5$ (cfr Section S4.1). For the Bases definition $\boldsymbol{O}_i$ is the geometrical center of the base beads, $\boldsymbol{O}_i = (\boldsymbol{B}_{1,i} + \boldsymbol{B}_{2,i})/2$. Finally, for the helical axis definition the local tangent vector $\hat{\boldsymbol{t}}_i$ was defined as the helical axis of the fragment including the base pairs $i-5, \ldots, i+5$

With these definitions, we computed the correlation functions $c_p, c_s$ and $c_d$ according to Equations (12), (13) and (14) in the main text. The resulting functions are plotted in Fig.S6. Extreme values and averages of the corresponding $l_p, l_s$ and $l_c$ are reported in Table S7.

| Definition | Statistical Feature | $l_p$ (nm) | $l_s$ (nm) | $l_d$ (nm) |
|---|---|---|---|---|
| | Average | 56 ± 1 | 326 ± 47 | 75 ± 3 |
| Sugars | Min | 46 | 101 | 62 |
| | Max | 64 | 786 | 117 |
| | Average | 63 ± 1 | 633 ± 108 | 77 ± 3 |
| Cylinder | Min | 54 | 94 | 69 |
| | Max | 72 | 1996 | 124 |
| | Average | 59 ± 1 | 460 ± 71 | 75 ± 3 |
| Bases | Min | 51 | 109 | 64 |
| | Max | 68 | 1302 | 109 |
| | Average | 60 ± 1 | 566 ± 87 | 74 ± 3 |
| Helical axis | Min | 49 | 83 | 66 |
| | Max | 69 | 1469 | 121 |

Table S7: Average, minimum and maximum values of $l_p, l_s$ and $l_d$ according to the various definitions.

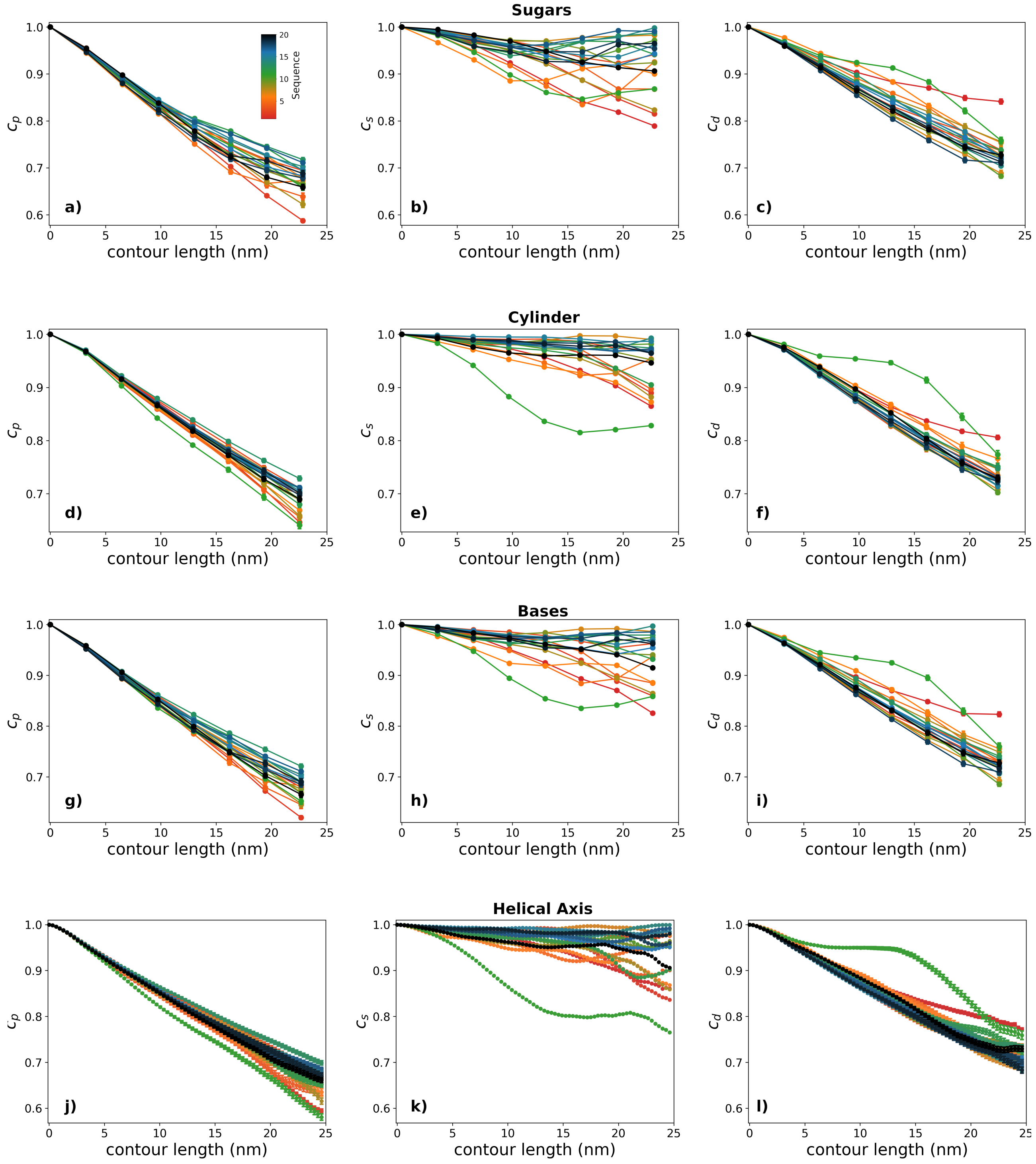


Figure S6: Correlation functions $c_p, c_s$ and $c_d$ obtained for the various sequences according to the four definitions. The sequences are listed in Section S3.2 and are numbered according to the colour code reported in panel a.

#### S7 Length dependence of elastic constants

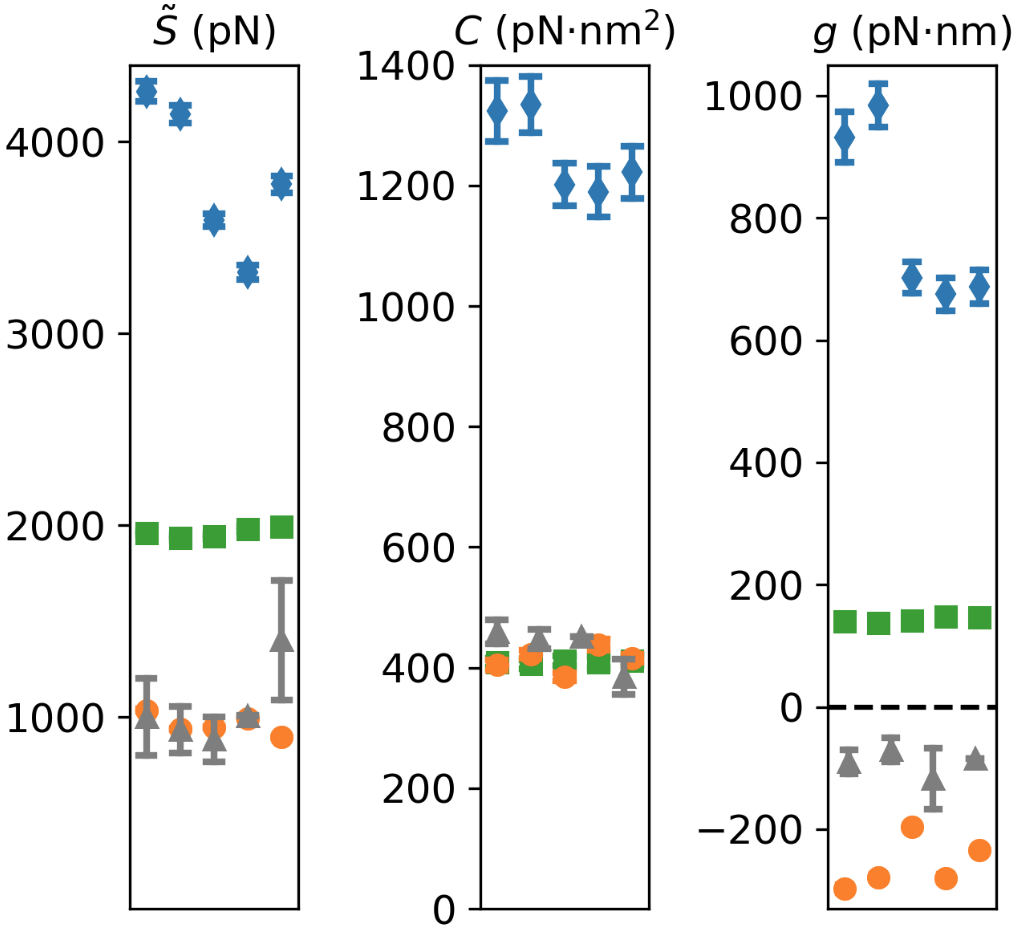


Figure S7: Elastic constants obtained for MADna (orange circles), oxDNA2 (green squares) and 3SPN2C (blue diamonds) for sequences made of twenty base pairs, which are listed as ST1-short,...,ST5-short in Section S3.3.

#### S8 Sequence-dependent helical pitch and persistence length for oxDNA2 and 3SPN2C

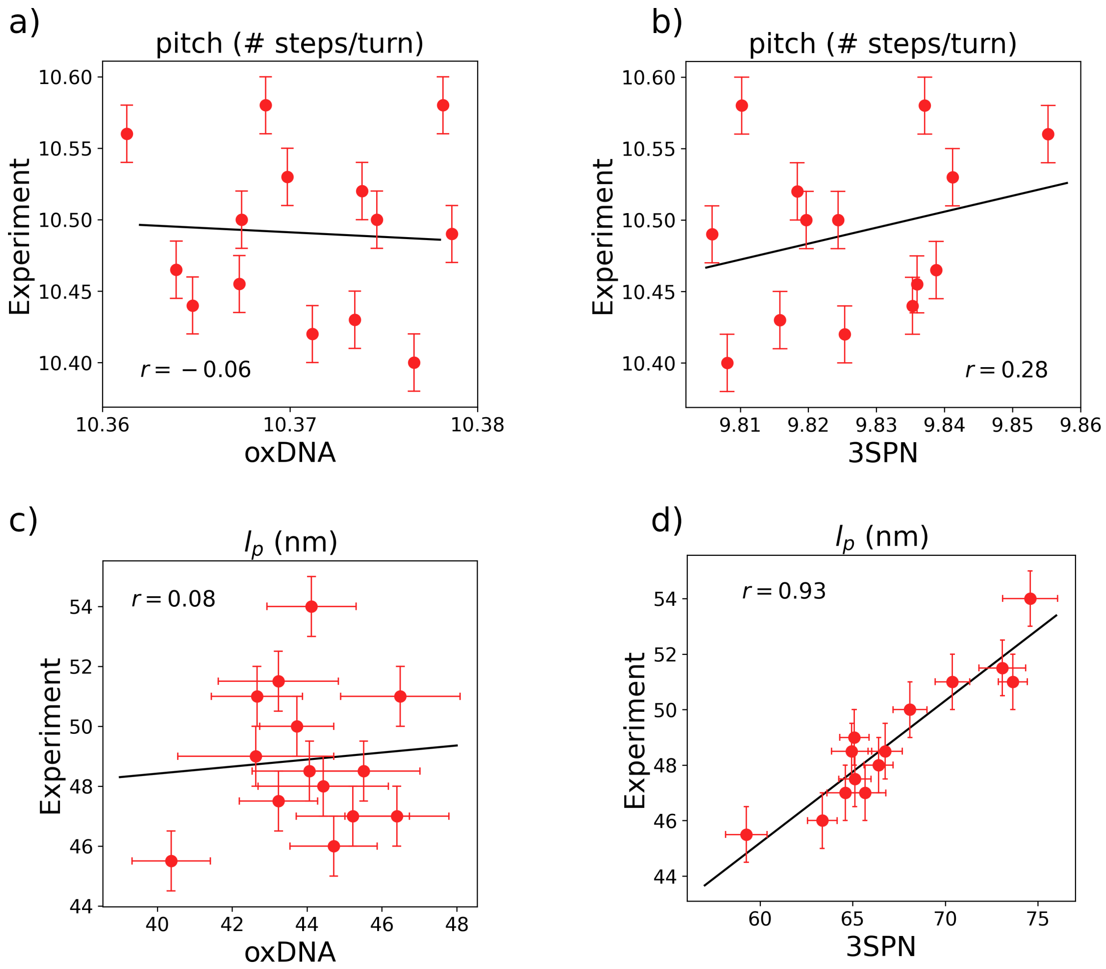


Figure S8: Comparison between experimental values and predictions by oxDNA2 and 3SPN2C for the sequence dependence of the helical pitch (a,b) and the persistence length $l_p$ (c,d). The lines correspond to the linear fits of the scatter plots and are included as a guide to the eye. The value of the Pearson coefficient is reported in each plot.

### S9 Convergence study of representative trajectories

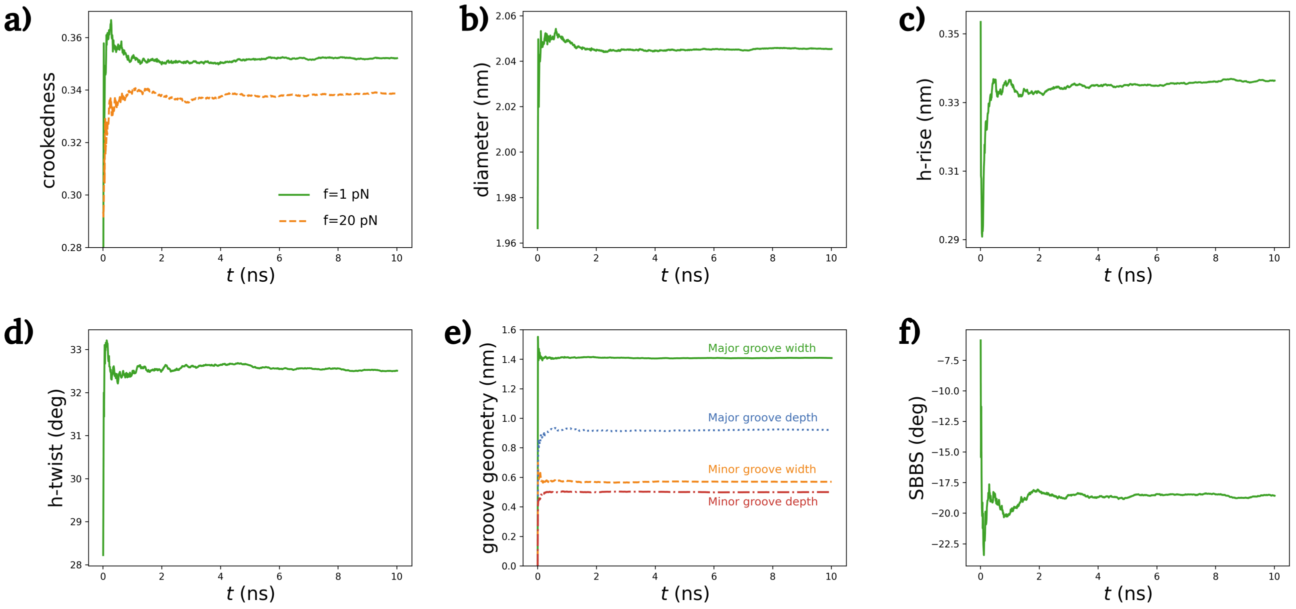


Figure S9: Convergence of the running average for various observables in a representative Benchmark simulation, corresponding to polyAA. In panels b)-f), we considered a simulation corresponding to $f = 1$ pN.

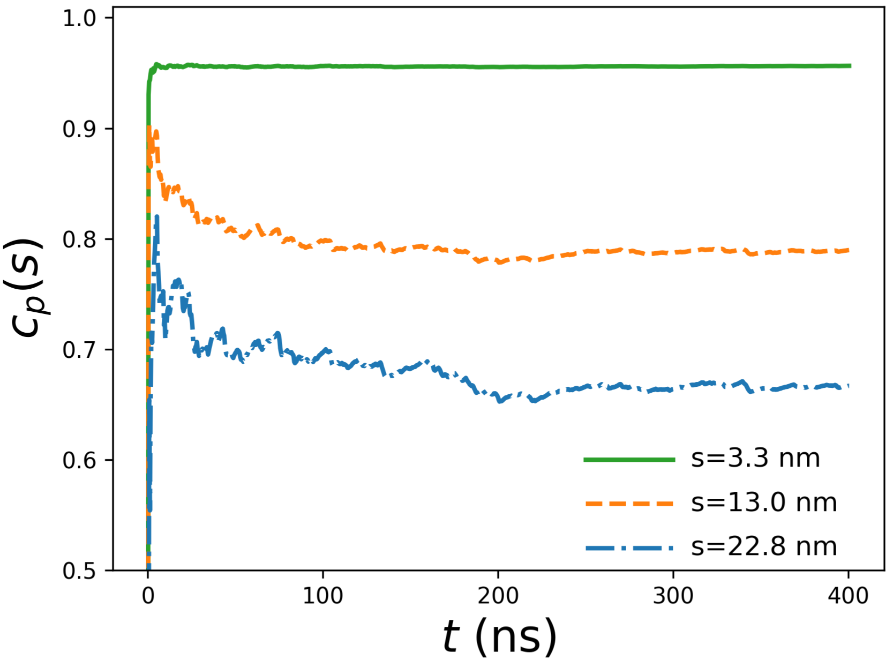


Figure S10: Convergence of the running average for the tangent vector correlation function $c_p(s)$ in a representative simulation for the computation of the sequence-averaged persistence length. We plot $c_p(s)$ for various values of the contour length $s$. The sequence considered in this representative case is P1.

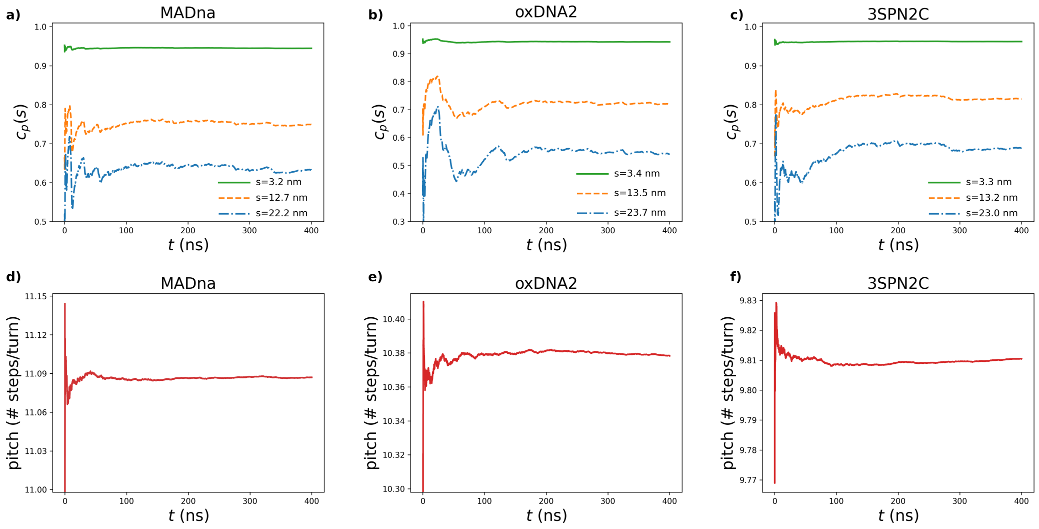


Figure S11: Convergence of the running average for the tangent vector correlation function $c_p(s)$ (top) and helical pitch (bottom) for a representative simulation for the computation of persistence length and helical pitch, for the three coarse-grained models. In the top panels, we plot $c_p(s)$ for various values of the contour length $s$. The sequence considered in this representative case is ACAT.

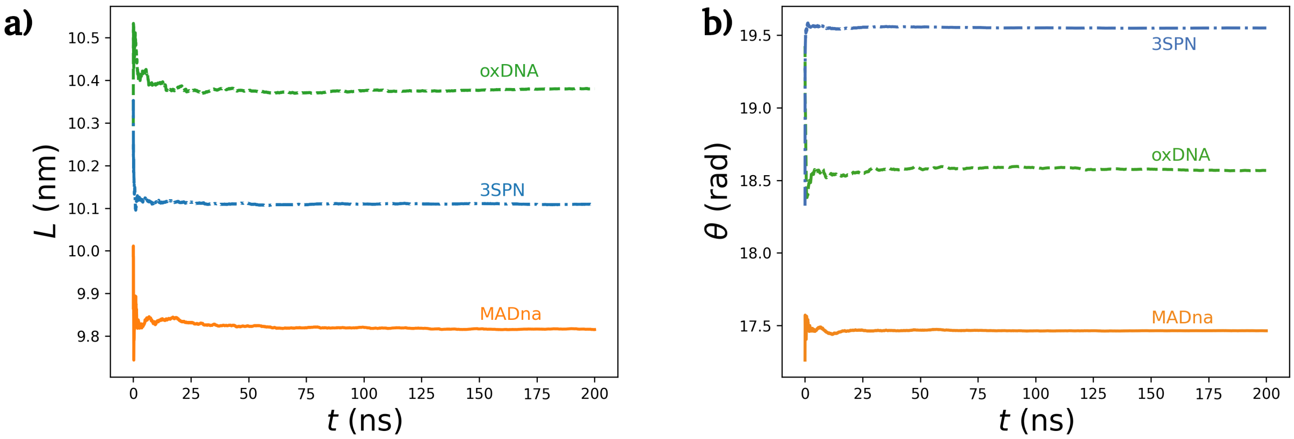


Figure S12: Convergence of the running average for the extension $L$ (a) and cumulate h-twist $\theta$ (b) for the three coarse-grained models, for a representative trajectory corresponding to sequence ST1 under the action of a force $f = 2$ pN and a torque $\tau = 5$ pN·nm.

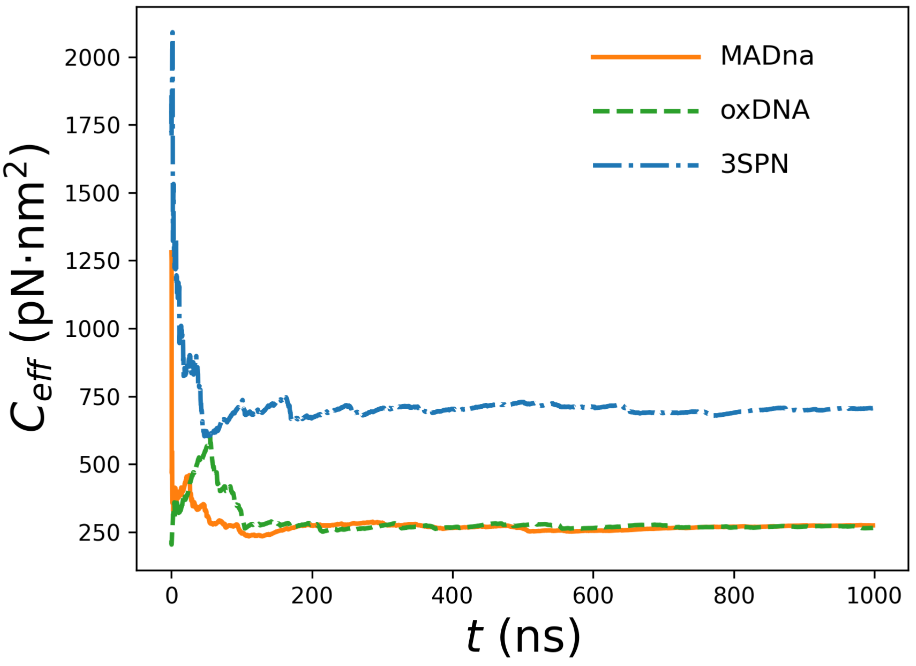


Figure S13: Convergence of the running average for effective twist modulus $C_{\text{eff}}$ in a representative simulation for the study of the twist-bend coupling for the three coarse-grained model. The representative trajectory corresponds to a simulation performed for sequence TB1 at a force $f = 0.75$ pN.

### Bibliography


[1] Hinckley, D. M., Freeman, G. S., Whitmer, J. K. and De Pablo, J. J. (2013) An experimentally-informed coarse-grained 3-site-per-nucleotide model of dna: Structure, thermodynamics, and dynamics of hybridization. *J. Chem. Phys.*, **139**, 144903.

[2] Marín-González, A., Vilhena, J. G., Moreno-Herrero, F. and Pérez, R. (2019) DNA Crookedness Regulates DNA Mechanical Properties at Short Length Scales. *Phys. Rev. Lett.*, **122**, 048102.

[3] Marín-González, A., Vilhena, J. G., Pérez, R. and Moreno-Herrero, F. (2017) Understanding the mechanical response of double-stranded DNA and RNA under constant stretching forces using all-atom molecular dynamics. *Proc. Nat. Academ. Sci.*, **114**, 7049–7054.

[4] Plimpton, S. (1995) Fast Parallel Algorithms for Short-Range Molecular Dynamics. *J. Comp. Phys.*, **117**, 1-19.

[5] Mitchell, J. S., Glowacki, J., Grandchamp, A. E., Manning, R. S. and Maddocks, J. H. (2017) Sequence-Dependent Persistence Lengths of DNA. *J. Chem. Theory Comput.*, **13**, 1539–1555.

[6] Eberly, D. (2020) Least Squares Fitting of Data by Linear or Quadratic Structures. www.geometrictools.com/Documentation/LeastSquaresFitting.pdf

[7] Wing, R., Drew., H., Takano, T., Broka, C., Tanaka, S., Itakura, K. and Dickerson, R. E. (1980) Crystal structure analysis of a complete turn of B-DNA *Nature*, **287**, 755–758.

[8] Lu, X.-J. and Olson, W. K. (2003) 3DNA: a software package for the analysis, rebuilding and visualization of three-dimensional nucleic acid structures. *Nucleic Acids Res.*, **31**, 5108–5121.
